# Novel ERR pan-agonists ameliorate heart failure through boosting cardiac fatty acid metabolism and mitochondrial function

**DOI:** 10.1101/2022.02.14.480431

**Authors:** Weiyi Xu, Cyrielle Billon, Hui Li, Matthew Hayes, Keyang Yu, McKenna Losby, Carissa S. Hampton, Christiana M. Adeyemi, Andrea Graves, Eleni Nasiotis, Chen Fu, Ryan Welch, Ronald M. Evans, Liming Pei, John K. Walker, Aleksandar Milosavljevic, Thomas Burris, Lilei Zhang

**Affiliations:** Department of Molecular & Human Genetics, Baylor College of Medicine, Houston, Texas, USA; Department of Pharmaceutical and Administrative Sciences, University of Health Sciences and Pharmacy, St. Louis, Missouri, USA; Center for Clinical Pharmacology, Washington University School of Medicine, St. Louis College of Pharmacy, St. Louis, Missouri, USA; Department of Pharmacology and Physiology, St. Louis University School of Medicine, St. Louis, Missouri, USA; University Hospitals Cleveland Medical Center, Cleveland, Ohio, USA; Howard Hughes Medical Institute, Salk Institute for Biological Studies, La Jolla, California, USA; Department of Pathology and Laboratory Medicine, University of Pennsylvania, Philadelphia, Pennsylvania, USA

## Abstract

Cardiac metabolic dysfunction is a hallmark of heart failure. Estrogen related receptors ERRα and ERRγ are essential regulators for cardiac metabolism. Therefore, activation of ERR could be a potential therapeutic intervention for heart failure. However, no natural or synthetic ERR agonist is available to demonstrate their pharmacological effect *in vivo*. Using a structure-based design approach, we designed and synthesized two structurally distinct pan-ERR agonists, SLU-PP-332 (332) and SLU-PP-915 (915), which significantly improved ejection fraction and ameliorated fibrosis against pressure overload-induced heart failure without affecting cardiac hypertrophy. Mechanistically, a broad-spectrum of metabolic genes were transcriptionally activated by ERR agonists, particularly genes involved in fatty acid metabolism and mitochondrial function, which were mainly mediated by ERRγ. Metabolomics analysis showed significant normalization of metabolic profiles in fatty acid/lipid and TCA/OXPHOS metabolites by 915 in the mouse heart with 6-week pressure overload. Autophagy was also induced by ERR agonists in cardiomycoyte. On the other hand, ERR agonism led to downregulation of cell cycle and development pathways, which was partially mediated by E2F1 in cardiomyocyte. In summary, ERR agonists maintain oxidative metabolism, which confers cardiac protection against pressure overload-induced heart failure *in vivo*. Our results provided direct pharmacological evidence supporting the further development of ERR agonists as novel heart failure therapeutics *in vivo*.

## Introduction

Heart failure is estimated to affect 6.5 million people in the US and 23 million people worldwide in 2020 (1). By 2030, the number of patients in the US is projected to be more than 8 million (2). Current therapeutic strategy for heart failure patients involves neurohormonal and sympathetic inhibition with the addition of device therapy (1, 3). The 5-year survival rate remains around 50% (4), suggesting an urgent need for novel therapy.

One of the hallmarks of heart failure is pathological metabolic remodeling where cardiac fuel metabolism and mitochondrial function are progressively dysregulated and impaired (5–7). Therefore, targeting cardiac metabolic regulators holds the potential to stop or even reverse the progression of heart failure (8). Previous studies in genetic mouse models demonstrated estrogen-related receptor (ERR) alpha and gamma target a large number of genes involved in regulating fatty acid and glucose metabolism, mitochondrial function, and muscle contraction in the heart (9–13). Moreover, transcriptomic analysis of human myocardium showed that a set of genes targeted by ERRα and its coactivator PGC1α were collectively downregulated in failing human hearts, and their expression levels correlated with the left ventricular ejection fraction (EF) (14). Therefore, ERRs have been proposed as candidate therapeutic targets for treating heart failure and other metabolic diseases (15, 16). Despite several ERR agonists having been previously developed (17–20), no study has demonstrated the potential utility for heart failure treatment *in vivo*. In this study, we report the cardiac protective effect of a recently described ERR pan-agonist SLU-PP-332 (332) and another novel structurally distinct ERR pan-agonist SLU-PP-915 (915) in pressure overload-induced heart failure. 332/915, mainly through ERRγ, activated a broad-specturm of metabolic genes and led to elevation in fatty acid metabolism and mitochondrial function, which improved the pumping function in TAC-induced heart failure *in vivo* without affecting cardiac hypertrophy.

## Methods

### Animals

All animal studies were approved by the Institutional Animal Care and Use Committee at Baylor College of Medicine and conducted in accordance with the NIH *Guide for the Care and Use of Laboratory Animals*. Mice were housed in a temperature- and humidity-controlled facility with a 12-h light/dark cycle and *ad libitum* access to water and standard laboratory rodent chow. 2- or 3-day-old Sprague-Dawley rat pups were purchased from Envigo and scarified for isolation of NRVMs upon arrival.

### Compound Design of SLU-PP-915

SLU-PP-915 was designed based on the ligand bound crystal structure of ERRγ with known acyl hydrazide agonist, GSK-4716 (PDB:2GPP) (21). Replacing the central hydrazide moiety with various 5-membered heterocycles led to a series of novel di-substituted thiophenes of ERRγ agonists, obtained in 2-steps from 5-bromo-2-thiophenecarboxylic acid.

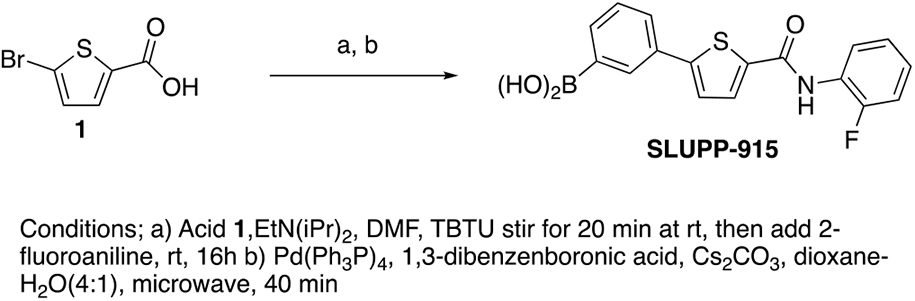

### Compound Synthesis of SLU-PP-915

1) 5-bromo-N-(2-fluorophenyl)thiophene-2-carboxamide

To a solution of 5-bromo-2-thiophenecarboxylic acid (0.100 g, 0.4830 mmol) and TBTU (0.1551 g, 0.4830 mmol) in dry DMF (4.8 mL) was added DIPEA (0.210 mL, 1.207 mmol). The reaction was stirred at room temperature for 20 min and then 2-fluoroaniline (0.0644 g, 0.5795 mmol) was added. The reaction was stirred overnight at room temperature. The reaction was quenched with water to precipitate the product. The mixture was filtered through a fritted funnel by vacuum suction. The solid was triturated with cold methanol, the solvent was removed by vacuum suction, and the product dried. The product was purified by flash chromatography (15% EtOAc:Hex) and isolated as a dark oil in 55%. ^1^H NMR (400 MHz, DMSO-d_6_) δ 10.30 (s, 1H), 7.91 (d, *J* = 4.0 Hz, 1H), 7.61 (t, *J* = 7.1 Hz, 1H), 7.43 (d, *J* = 4.0 Hz, 1H), 7.38-7.32 (m, 2H), 7.30-7.25 (m, 1H), 2.74 (s, 1H). LCMS: m/z calcd. for C_11_H_17_BrFNOS [M+H]^+^: 299.9 & 301.9; found [M+H]^+^: 299.9 & 302.0.

2) (3-(5-((2-fluorophenyl)carbamoyl)thiophen-2-yl)phenyl)boronic acid (SLU-PP-915) To a microwave reactor vessel was added sequentially 1,3-benzenediboronic acid (0.1143 mg, 0.86 mmol), Tetrakis(triphenylphosphine)palladium(0) (0.077 g, 0.66 mmol), Cs_2_CO_3_ (0.470 g, 1.33 mmol) and the above thiophene bromide (0.200 g, 0.66 mmol). The mixture was dissolved in a 4:1 mixture of dioxane: H_2_O (10 mL) and the vessel was sealed. The reaction was heated to 120°C for 40 min in the microwave and cooled to RT. The reaction mixture was washed with EtOAc through pad of celite and evaporated to afford a black solid. The crude product was purified by high pressure reverse phase column. The combined fractions were dried, sonicated with hexane and then washed with DCM several times and then filtered under gravity to afford SLU-PP-915 (0.061 mg, 53.8%) as a white solid.^1^H NMR (400 MHz, DMSO-d_6_) δ 10.17 (s, 1H), 8.23 (s, 2H), 8.17 (s, 1H), 8.03 (d, *J* = 4.0 Hz, 1H), 7.84-7.77 (m, 2H), 7.62-7.59 (m, 2H), 7.44 (t, *J* = 7.6 Hz, 1H), 7.34-7.27 (m, 2H), 7.27-7.21 (m, 1H); HRMS m/z calcd for (C_17_H_13_BFNO_3_S)-H 340.0620, found 340.0620.

### Co-transfection assays

HEK293 cells were maintained in Dulbecco’s modified Eagle’s medium (DMEM) supplemented with 10% fetal bovine serum at 37 °C under 5% CO_2_. Twenty-four hours prior to transfection, HEK293 cells were plated in 96-well plates at a density of 2×10^4^ cells/well. GAL4-NR-LBD, or FLAG-ERR-FL plasmids were used in the luciferase assay.

### Pharmacokinetic studies of SLU-PP-915

Pharmacokinetic studies of SLU-PP-915 in mice were performed as previously described (22). Three-month old C57Bl6/J male mice (n=4) were intraperitoneally injected at ZT0 with 20mg/kg of SLU-PP-915 (10% Cremophor-10% DMSO-80% PBS). Animals were sacrificed by CO_2_ asphyxiation and tissues were collected at 1h, 2h or 4h after administration of the compound (n=4 per time point). Plasma were collected and flash frozen and stored at −80 °C until analysis. Plasma samples were analysed by Charles River.

### Preparation and administration of SLU-PP-332 and SLU-PP-915

For *in vitro* study, both compounds were prepared in 10mM in DMSO as stock and used at 10μM in cell culture medium. Stock solutions were aliquoted and stored at −20°C. For *in vivo* study, 332 and 915 were prepared in 12% DMSO, 15% Cremophor in PBS without Ca^2+^/Mg^2+^ at 5mg/ml, and intraperitoneally administered to mice twice a day at ZT0 and ZT12 at the dose of 25mg/kg. Powder was first dissolved in DMSO as stock which was aliquoted and stored at −20°C. Aliquot of stock in DMSO was further diluted with Cremophor with gentle shaking and stored at 4°C for no more than 5 days. Immediately before use, solution was diluted in PBS without further storage.

### Transaortic constricution (TAC)

TAC was performed as previously described (23). Briefly, a skin incision was made from the neck to sternum, and the muscle and soft tissues were dissected to expose trachea. The intubation tube was gently inserted until the tube was seen inside the trachea. The endotracheal tube was then connected to a Harvard volume-cycled rodent ventilator cycling at 125-150 breaths/minute and a tidal volume of 0.1-0.3ml. The sternum was cut in the middle, and the aortic arch was exposed under a surgical microscope. The aortic arch was separated and freed between the right innominate artery and left carotid artery, and was then looped by a 7-0 silk suture. A blunted 27-gauge needle was placed on top of the aorta, and the silk suture was tied around the needle. The needle was then carefully removed from under the suture. The body wall and skin were closed with 5-0 nylon suture separately. Sham operated animals underwent a similar procedure without constriction of the aorta.

### Echocardiography

Echocardiography was performed as previously described (23). Briefly, mice were anesthetized by 1% isoflurane supplied with 100% oxygen through the nose cone and placed in a temperature-controlled (40-42°C) ECHO stage to keep the body temperature range from 36.5-37.5°C. Warm echo gel was placed on the shaved chest and echography was taken using a 32-56 MHz probe in the parasternal short axis at the papillary muscle level. Images were taken using Vevo 2100 High Resolution Imaging System (Visual Sonics Inc.) under M-mode and analyzed using ECHO workstation. At least three beats were measured and averaged for analysis.

### Histological staining

Masson trichrome staining for fibrosis were performed as previously described (23). Briefly, short-axis heart sections obtained from the midventricular area were fixed in PBS with 4% paraformaldehyde and embedded in paraffin. Sections were then stained using Trichrome stain (Masson) kit (Sigma-Aldrich, HT15) for differential staining of collagen (blue) and muscle fiber (red). Sections were then imaged and fibrotic area was quantified by ImagePro software. Cardiomyocyte cross-sectional area was assessed by staining with wheat germ agglutinin (WGA) conjugated with Alexa Fluor 594 (Thermo Fisher Scientific, W11262). Sections were imaged and individual cardiomyocyte size was quantified using ImageJ software (NIH). In each cross-section, 90-330 cells were counted for quantification.

### Transmission electron microscopy

Cardiac muscle tissue obtained from LV free wall were sent to the electron microscopy core at Washington University in Saint Louis for processing and imaging. Ultrastructural heart morphology was examined using transmission electron microscopy. Cardiac muscle tissue blocks were fixed in 2.5% glutaraldehyde in 0.1 M phosphate buffer (pH=7.4). Post-fixation was performed in 1% OsO_4_ in 0.1 M cacodylate buffer (pH=7.4) supplemented with 1.5% K_4_[Fe(CN)_6_]. Subsequently, samples were dehydrated and embedded in epon. Ultrathin sections were examined using a JEOL 1200EX electron microscope.

### Neonatal rat ventricular myocytes (NRVMs)

NRVMs were isolated and cultured as previously described (23). Briefly, freshly isolated hearts from rat pups were rinsed in Hanks’ Balanced Salt Solution (HBSS) and incubated in 0.05% trypsin EDTA at 4°C overnight. On the next day, cardiomyocytes were dissociated with type 2 collagenase (Worthington, LS004176) in HBSS and neutralized with 10% FBS in DMEM. The cell suspension was further incubated in a tissue culture-treated plastic dish for 90 min to allow fibroblast attachment. Typically, NRVMs were seeded at 50-70% confluence in DMEM supplemented with 10% FBS, 1% penicillin/streptomycin and 100μM bromodeoxyuridine. 2 days after seeding, the medium was changed to DMEM supplemented with insulin/transferrin/selenium (Sigma-Aldrich, I3146) and cultures were used within 5 days. To induce hypertrophy, NRVMs were treated with 100μM phenylephrine (PE) for 48hr. For siRNA knockdown, NRVMs were transfected with siRNA against rat *Esrra* (Dharmacon, M-085000-02-0005), rat *Esrrb* (Dharmacon, M-100126-01-0005), rat *Esrrg* (Dharmacon, M-100212-01-0005), rat *E2f1* (Dharmacon, M-102754-00-0005) or scramble control siRNA (ThermoFisher, 4390843), using the Viromer BLUE Transfection Reagent (Lipocalyx, VB-01LB-01). Typically, 25 nM siRNA was used for NRVMs seeded in 24-well.

### Measurement of oxygen consumption rate

Adult mouse cardiomyocytes were isolated by Langend orff perfusion and cultured as previously described (24). Isolated mouse cardiomyocytes were seeded at 2K per well in the Seahorse XF96 cell culture microplates (Agilent Technologies, 102416100) pre-coated with laminin mouse protein at 2μg/cm^2^ (Thermo Fisher Scientific, 23017015) and maintained in the culture medium with 2% CO_2_ (24). Cells were treated with vehicle or 915 for 24hr. Myosin inhibitor 2,3-butanedione monoxime (BDM) (Sigma-Aldrich, B0753) was added in the culture medium to improve viability and rod-shape morphology (24, 25). Oxygen consumption rate was measured in extracellular flux analyzer XFe96 (Agilent) with pyruvate (Sigma-Aldrich, P2256) or palmitate-BSA conjugate (Cayman Chemical, 29558) as substrate following the manufacturer’s protocol and previous publication (26). 1.5 μM oligomycin (Sigma-Aldrich, 495455), 1 μM FCCP (Sigma-Aldrich, C2920), 5μM rotenone (Sigma-Aldrich, 45656), and 5μM antimycin A (Sigma-Aldrich, A8674) were used for the assay protocol (27). The maximum mitochondria respiratory capacity was measured as the difference between plateau value after injecting FCCP and the end-point value after injecting rotenone and antimycin A. OCR was normalized to DNA content in each well measured by CyQUANT® Cell Proliferation Assay kit (Thermo Fisher Scientific, C7026) (28).

### RNA and DNA extraction

For RT-PCR, RNA from cell culture and mouse heart tissue was extracted using high pure RNA isolation kit (Roche, 11828665001) and miRNeasy Mini kit (Qiagen, 217004), respectively. For RNA-Seq, RNA from NRVMs was extracted using miRNeasy Mini kit (Qiagen, 217004).

DNA from mouse heart ventricular tissues or NRVMs was extracted using QIAamp® DNA Mini Kit (Qiagen, 51304) according to the manufacture’s procotol. Mouse heart tissues were homoginated with stainless steel beads before DNA extraction.

### RT-PCR

Reverse transcription was performed using iScript™ Reverse Transcription Supermix (Bio-Rad Laboratories, 1708841). Real-time Taqman-PCR was assembled in qPCRBIO Probe Blue Mix (Genesee Scientific Corporation, 17-514) or TaqMan Fast Advanced Master Mix (Thermo Fisher Scientific, 4444554) with 0.5μM of each primer (Integrated DNA Technologies) and 50nM Universal ProbeLibrary Probe (Roche). RT-PCR was performed in QuantStudio™ 5 Real-Time PCR System (Thermo Fisher Scientific) and calculated using ^ΔΔ^Ct-method. Expression level of *Ppib* was used as internal control unless otherwise indicated.

For quantification of mitochondrial DNA copy number, the RT-PCR reaction was assembled in SsoAdvanced™ Universal SYBR® Green Supermix (Bio-Rad, 1725271) with 0.5μM of each primer. Expression level of *Chr4* (rat chromosome 4) and *Chr6* (mouse chromosome 6) was used as internal control for NRVMs and mouse heart tissues (29), respectively. Primer list is available upon request.

### RNA-Seq and analysis

Isolated RNA from NRVMs was sent to Genome Technology Access Center (GTAC), Washington University in Saint Louis for sequencing. mRNA were purified using PolyA selection. Libraries with uniquely dual-indexed with 10bp fragments were generated from mRNA. Sample libraries were multiplexed onto the appropriate amount of Illumina NovaSeq S4, XP 2×150 runs. 30 million clusters were used for gene expression profiling studies. RNA-seq results were analyzed using GTAC@MGI standard RNA-seq analytical pipeline. This comprehensive analysis of RNA-seq data includes multiple levels of quality control, differential expression, and pathway analysis.

Differentially expressed genes were identified as genes with an adjusted p-value of less than 0.05 and fold change greater than 1.5 between the control and treatment group. Gene ontology and pathway enrichment analysis were performed using ShinyGO v0.61 bioinformatics suite (30). An area-proportional Venn diagram was created using BioVenn (31). Motif analysis was performed with HOMER.

Isolated RNA from mouse heart tissue was sent to the Salk Institute for RNA sequenqcing and data analysis was performed at Baylor using deconvoluation described below.

### Deconvolution of bulk RNA-seq data of mouse heart

1) XDec deconvolution algorithm

XDec is a reference-free cell type deconvolution algorithm that predicts the proportion of and the expression profile of constituent cell types of complex samples using RNAseq data [Murillo et al., 2022, paper under review]. Three stages of XDec, stage 0, stage 1, and stage 2. Briefly, informative features that separates known cell types of the complex sample are identified from single-cell RNA expression profile of known cell types in stage 0. The XDec then performs deconvolution through constrained matrix factorization using the selected features and bulk-RNAseq profile of complex sample in stage 1 and 2.

2) scRNA sequencing data processing

Single cell RNA sequencing data of known major cell types of mice heart with TAC treatment was downloaded from GEO (GSE120064). Metadata of the same study were used to identify the cell type of each single cell expression profile, including cardiomyocyte, fibroblast, endothelial cell, macrophage, granulocyte, and T cells (32). Since there is a limited proportion of immune cells in the mice heart, the granulocyte, macrophage, and T cell profiles were grouped to represent immune cells. Profiles are ranked within each cell types based on coverage and those with the lowest 25 percentile of coverage were excluded from the study. Pseudo-bulk profiles of each cell type were then created by performing addition across all genes for every 10 profiles (33). For each of the four cell types, cardiomyocyte, fibroblast, endothelial cell, and immune cell, 100 pseudo-bulks were selected to determine informative features in XDec stage 0. Expression count of the pseudo-bulk profiles were quantile normalized and then transformed into the constrained [0,1] range. The transformed profile was then filtered for genes that were present in 332 bulk RNAseq data to ensure features selected by XDec stage 0 would inform deconvolution.

3) Processing of bulk RNAseq data from experimental samples for XDec

Expression profile of 332 and vehicle treated TAC and Sham mice heart derived from bulk RNAseq sequencing files were combined into a gene expression matrix. All samples were quantile normalized. The resulting expression matrix was the untransformed expression profile to be used in XDec stage 2. The matrix is then transformed into constrained values in the [0,1] range for stage 1 XDec deconvolution. Genes that are not expressed in any of the samples or are not present in the single cell RNA seq profiles of TAC mice (GSE120064) were removed from the transformed matrix.

4) TAC mice heart XDec feature selection

Informative features were selected based on XDec stage 0 performed on transformed pseudo-bulk expression profiles of cardiomyocyte (CM), fibroblast (FB), endothelial cell (EC), and immune cell (IM) in TAC mice heart derived from dataset GSE120064. T-tests were performed to compare transformed gene expression counts between pairs of cell types and between one cell type and the rest. Top up- and down-regulated genes that are significantly different based on t-tests were selected as informative features.

To identify profiles that are unique to 332 treatments, we also randomly selected 15 differentially expressed genes between 332 treated and vehicle treated TAC mice heart bulk RNAseq profiles to add to the informative features. This produced a list of 184 unique informative features that would guide XDec deconvolution (Table 2).

**Table 1.** DEGs identified as ERRα/γ target genes in previous study.

| Name | 332 DEGs | 915 DEGs | ERRα binding | ERRγ binding |
| --- | --- | --- | --- | --- |
| *Pdk4* | x | x |  | x |
| *Acadm* | x | x | x | x |
| *Hk2* | x | x | x |  |
| *Ndufb5* | x | x | x | x |
| *Fabp3* | x | x |  | x |
| *Cox8b* | x | x |  | x |
| *Cyp27a1* | x | x |  | x |
| *Etfb* | x | x |  | x |
| *Myl3* | x | x | x | x |
| *Atp5g1* | x |  |  | x |
| *Ppif* | x |  |  | x |
| *Esrra* | x |  | x | x |
| *Csmd1* | x |  |  | x |
| *Cycs* | x |  | x | x |
| *Mrps18b* |  | x | x |  |
| *Mrpl47* |  | x | x | x |
| *Ldhb* |  | x | x | x |
| *Me3* |  | x |  | x |
| *Ndufs7* |  | x | x | x |
| *Bdnf* |  | x | x | x |
| *Cenpt* | x | x | x | x |
| *Ppl* | x | x |  | x |
| *Lmln* | x |  | x |  |
| *Fgl2* | x |  | x | x |
| *Cbs* | x |  | x |  |
| *Abhd8* |  | x |  | x |
| *Ets1* |  | x |  | x |
| *Krt8* |  | x |  | x |
| *Vegfc* |  | x | x |  |
| *Htr2b* |  | x | x |  |
| *Pi15* |  | x | x |  |
| *Cenpa* |  | x | x | x |

Up-regulated and down-regulated DEGs are labeled in red and blue, respectively.

**Table 2.** List of feature genes used for deconvolution of bulk RNA-seq data.

List of feature genes used for deconvolution of bulk RNA-seq data
|  |  |  |  |  |  |
| --- | --- | --- | --- | --- | --- |
| ENSMUSG00000019787 | ENSMUSG00000030116 | ENSMUSG00000032942 | ENSMUSG00000041445 | ENSMUSG00000037813 | ENSMUSG00000008035 |
| ENSMUSG00000032366 | ENSMUSG00000023224 | ENSMUSG00000081964 | ENSMUSG00000033082 | ENSMUSG00000052374 | ENSMUSG00000055116 |
| ENSMUSG00000019505 | ENSMUSG00000026879 | ENSMUSG00000010122 | ENSMUSG00000074874 | ENSMUSG00000040752 | ENSMUSG00000037519 |
| ENSMUSG00000092341 | ENSMUSG00000074218 | ENSMUSG00000029685 | ENSMUSG00000001930 | ENSMUSG00000068614 | ENSMUSG00000053646 |
| ENSMUSG00000060591 | ENSMUSG00000035458 | ENSMUSG00000090889 | ENSMUSG00000032878 | ENSMUSG00000028116 | ENSMUSG00000039220 |
| ENSMUSG00000001175 | ENSMUSG00000056366 | ENSMUSG00000049709 | ENSMUSG00000044921 | ENSMUSG00000054889 | ENSMUSG00000045776 |
| ENSMUSG00000078193 | ENSMUSG00000091898 | ENSMUSG00000059852 | ENSMUSG00000045930 | ENSMUSG00000024049 | ENSMUSG00000009633 |
| ENSMUSG00000091449 | ENSMUSG00000025488 | ENSMUSG00000039395 | ENSMUSG00000032232 | ENSMUSG00000059447 | ENSMUSG00000072902 |
| ENSMUSG00000090733 | ENSMUSG00000000682 | ENSMUSG00000005220 | ENSMUSG00000024168 | ENSMUSG00000038239 | ENSMUSG00000038319 |
| ENSMUSG00000036438 | ENSMUSG00000069792 | ENSMUSG00000023019 | ENSMUSG00000020262 | ENSMUSG00000029467 | ENSMUSG00000078921 |
| ENSMUSG00000040952 | ENSMUSG00000006519 | ENSMUSG00000055027 | ENSMUSG00000068079 | ENSMUSG00000031461 | ENSMUSG00000011256 |
| ENSMUSG00000069439 | ENSMUSG00000060063 | ENSMUSG00000044951 | ENSMUSG00000034911 | ENSMUSG00000067653 | ENSMUSG00000030089 |
| ENSMUSG00000031871 | ENSMUSG00000039109 | ENSMUSG00000085277 | ENSMUSG00000086584 | ENSMUSG00000021798 | ENSMUSG00000020032 |
| ENSMUSG00000022579 | ENSMUSG00000018930 | ENSMUSG00000063428 | ENSMUSG00000066842 | ENSMUSG00000053093 | ENSMUSG00000004098 |
| ENSMUSG00000002944 | ENSMUSG00000036896 | ENSMUSG00000044716 | ENSMUSG00000071001 | ENSMUSG00000021313 | ENSMUSG00000057137 |
| ENSMUSG00000079018 | ENSMUSG00000058715 | ENSMUSG00000026824 | ENSMUSG00000022206 | ENSMUSG00000047419 | ENSMUSG00000024560 |
| ENSMUSG00000026921 | ENSMUSG00000036905 | ENSMUSG00000005628 | ENSMUSG00000062329 | ENSMUSG00000029019 | ENSMUSG00000034641 |
| ENSMUSG00000020577 | ENSMUSG00000030579 | ENSMUSG00000025221 | ENSMUSG00000002020 | ENSMUSG00000051747 | ENSMUSG00000024487 |
| ENSMUSG00000022661 | ENSMUSG00000018593 | ENSMUSG00000035296 | ENSMUSG00000037379 | ENSMUSG00000031698 | ENSMUSG00000022307 |
| ENSMUSG00000022584 | ENSMUSG00000006356 | ENSMUSG00000030621 | ENSMUSG00000032625 | ENSMUSG00000059741 | ENSMUSG00000023046 |
| ENSMUSG00000022018 | ENSMUSG00000029309 | ENSMUSG00000029651 | ENSMUSG00000052957 | ENSMUSG00000018893 | ENSMUSG00000022665 |
| ENSMUSG00000062515 | ENSMUSG00000020315 | ENSMUSG00000039376 | ENSMUSG00000032334 | ENSMUSG00000028773 | ENSMUSG00000063904 |
| ENSMUSG00000093490 | ENSMUSG00000031502 | ENSMUSG00000021373 | ENSMUSG00000055632 | ENSMUSG00000030785 | ENSMUSG00000050796 |
| ENSMUSG00000025190 | ENSMUSG00000064349 | ENSMUSG00000024471 | ENSMUSG00000000567 | ENSMUSG00000038583 | ENSMUSG00000059713 |
| ENSMUSG00000079017 | ENSMUSG00000016494 | ENSMUSG00000010175 | ENSMUSG00000031595 | ENSMUSG00000026414 | ENSMUSG00000008845 |
| ENSMUSG00000000628 | ENSMUSG00000037712 | ENSMUSG00000046603 | ENSMUSG00000007888 | ENSMUSG00000013936 | ENSMUSG00000028955 |
| ENSMUSG00000026574 | ENSMUSG00000008575 | ENSMUSG00000042448 | ENSMUSG00000031343 | ENSMUSG00000095150 | ENSMUSG00000038762 |
| ENSMUSG00000000753 | ENSMUSG00000020542 | ENSMUSG00000044258 | ENSMUSG00000043613 | ENSMUSG00000060802 | ENSMUSG00000032267 |
| ENSMUSG00000021876 | ENSMUSG00000037905 | ENSMUSG00000054385 | ENSMUSG00000071005 | ENSMUSG00000061848 | ENSMUSG00000022793 |
| ENSMUSG00000036446 | ENSMUSG00000021898 | ENSMUSG00000052921 | ENSMUSG00000027797 | ENSMUSG00000037706 |  |
| ENSMUSG00000029718 | ENSMUSG00000003476 | ENSMUSG00000028996 | ENSMUSG00000060572 | ENSMUSG00000060467 |  |

5) Application of XDec deconvolution to bulk RNAseq data

Since the number of bulk-RNAseq profiles from the mouse heart samples is limited (n=12), we included other TAC mice heart profiles (n=17) and two reference TAC mice heart profiles for each known cell type (GSE120064) to improve accuracy.

XDec performed stage 1 on the transformed expression matrix of the combined samples using the 184 informative features identified in stage 0. XDec stability test was performed to identify optimal number of constituent cell types within a range of [4,16]. Based on the stability test result, XDec deconvolution was performed with a parameter of 11 constituent cell types, max iteration of 2000 and a maximum difference of residual sum of squares between consecutive iterations of 1e-10.

We performed correlation test between all transformed reference profiles and transformed estimated expression profile of the 11 constituent cell types over the informative features to discern cell type identity. Since multiple CM profiles were identified, estimated proportion of the 11 constituent cell types in each sample is then compared to identify treatment specific profiles. The untransformed estimated expression profiles of the 11 constituent cell types were then obtained by performing XDec stage 2 on the estimated proportion and the untransformed expression matrix of all the mouse heart samples in the experiment.

6) Differential expression analysis of XDec estimated profiles

We performed t-test adjusted for average comparison on the average untransformed gene expression value of all genes between SLU 332 treated TAC CM profile and vehicle treated TAC CM profile. P-value is adjusted by FDR. Genes with an FDR adjusted p-value < 0.05 and a t statistic larger than 0 were identified as differentially expressed genes (DEGs).

7) Pathway enrichment analysis

KEGG pathway enrichment analysis was performed on the up-regulated DEGs and down-regulated DEGs separately. The analysis is performed using KEGGREST package in R (34) and KEGG PATHWAY database (35). Significantly enriched pathways were called based on p-value < 0.05.

### Immunoblotting

NRVMs were lysed in RIPA buffer (Thermo Fisher Scientific, 89901) supplemented with protease inhibitor cocktail (Roche, 11836153001). Protein concentration was determined using BCA assay kit (Thermo Fisher Scientific, 23225). Protein samples were mixed with LDS sample buffer (Thermo Fisher Scientific, B0008) and reducing agent (Thermo Fisher Scientific, B0009), and then incubated at 95°C for 10min. Proteins were separated on 4-12% Bis-Tris gels (Thermo Fisher Scientific) with MES SDS running buffer (Thermo Fisher Scientific, B0002). For immunoblotting of LC3B, proteins were separated on 16% tricine gels (Thermo Fisher Scientific, EC6695BOX) with tricine SDS running buffer (Thermo Fisher Scientific, LC1675). Protein was then transferred to 0.2μm PVDF membrane (Bio-rad, 1704272) by semi-dry transfer at 25V for 30min using Trans-Blot^®^ Turbo™ Transfer System (Bio-rad). PVDF membrane was then blocked with 5% nonfat milk (Bio-rad, 1706404) in tris buffered saline with 0.1% Tween (TBST) for 1hr at room temperature. Membrane was then incubated with primary antibody at 4°C overnight and horseradish peroxidase (HRP)-conjugated secondary antibody at room temperature for 1hr. Enhanced chemiluminescence substrate (Thermo Fisher Scientific, 34580 or 34094) of HRP was used for detection. Band intensity was quantified by ImageJ (NIH) and GAPDH was used as internal control. Primary antibodies used are listed as below: anti-phosph-ERK1/2 (Cell Signaling Technologies, 9101S), anti-ERK1/2 (Cell Signaling Technologies, 9102S), anti-LC3B (Cell Signaling Technologies, 2775S), anti-p62 (Cell Signaling Technologies, 8025S or 5114S), anti-Vinculin (Cell Signaling Technologies, 18799S), anti-ERRα (Abcam, ab76228), anti-GAPDH (Sigma-Aldrich, G8795) and anti-E2F1 (Sigma-Aldrich, SAB2103144). All were used at 1:1000 dilution. Primary antibody for ERRγ has been described (10).

### Metabolomics

Metabolite analysis of cardiac muscle tissue obtained from LV free wall was conducted by Metabolon Inc. Samples with low quality or high degradation were excluded from analysis.

### Luciferase reporter assay

Transcriptional activity of NFAT was measured in NRVM as previously described (36, 37). Briefly, NRVM was infected with adenovirus containing NFAT-responsive luciferase reporter construct (Welgen Inc., S3010) at the multiplicity of infection of 400. Treatment on NRVMs was performed 24hr post-infection, and luciferase assay was performed by using dual-Luciferase reporter assay kit (Promega, E1910). Emission of light was measured in FLUOstar Omega microplate reader and analyzed with the built-in software (BMG LABTECH). Luminesence intensity was normalized to protein content in each well.

## Results

### ERR agonists improved cardiac function against TAC-induced heart failure in vivo

To induce heart failure *in vivo*, we performed transverse aortic constriction (TAC) in mice, a classic model for pressure overload-induced cardiac hypertrophy and heart failure (38). Echocardiogram was perform to assess cardiac function (Figure 1A, 2A-B and Supplementary Figure 1A-I). A reduction in cardiac output measured by ejection fraction (EF) was first observed at 2 weeks post TAC, which further decreased to 33.8% compared to 68.2% of the sham group at 6 weeks (Figure 1A). To study the pharmacological effects of ERR activation on pressure overload-induced heart failure, we started with the previously described ERR pan-agonist 332 and administered the drug to mice post TAC surgery by intraperitoneal injection twice daily throughout the 6-week experiment. Our results showed that 332 significantly improved the EF after TAC at 4-6 weeks (Figure 1A). 332 also significantly reduced the expression level of cardiac stress-induced genes *Nppa* and *Nppb* (Figure 1B). Furthermore, the Masson’s trichrome staining demonstrated an increase in cardiac fibrosis after TAC, which is diminished by 332 treatment (Figure 1C). Together, these data show that 332 improved pump function and relieved the cardiac stress and subsequent injury and fibrosis induced by pressure overload.

**Figure 1.**
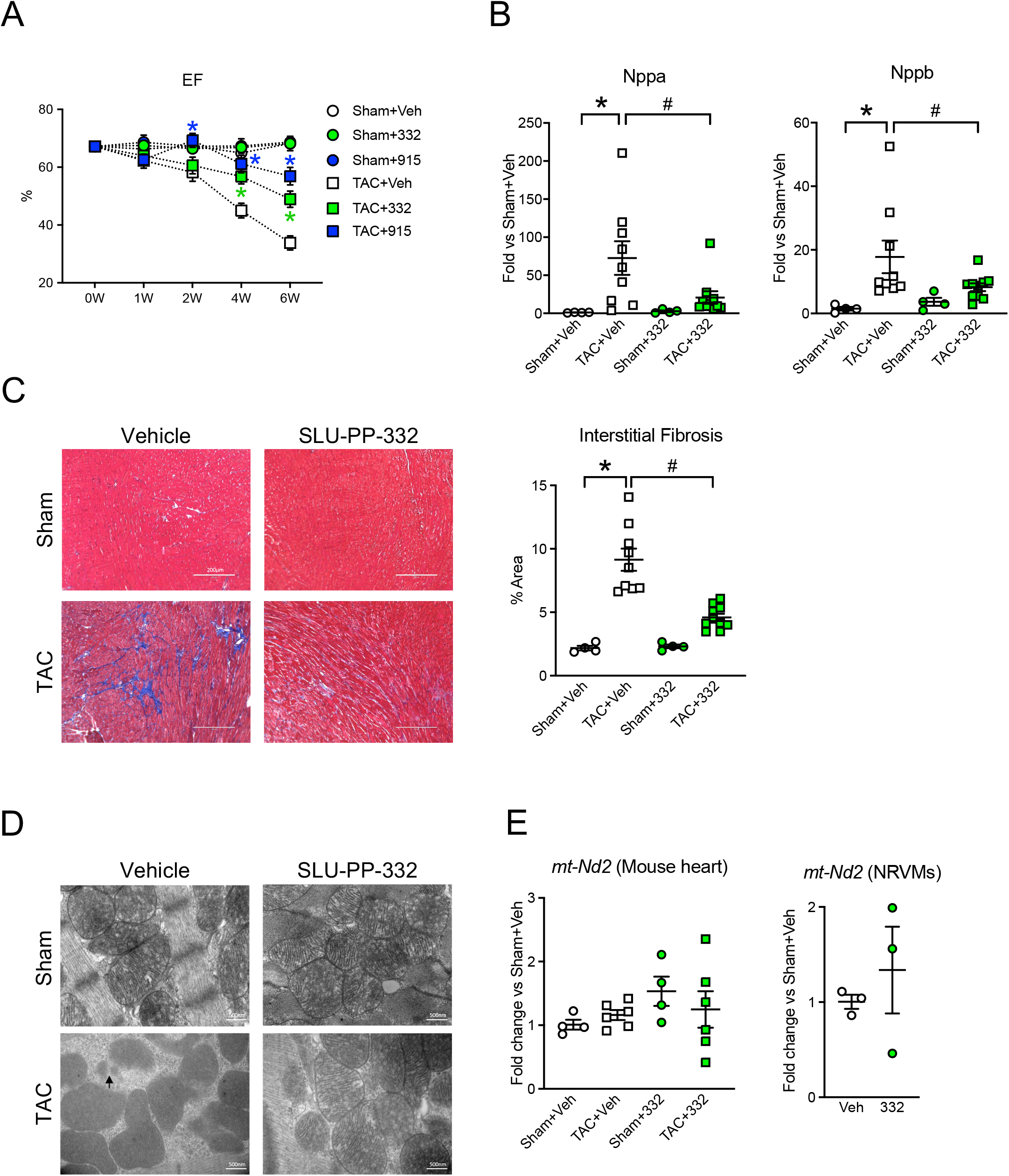
ERR agonists attenuate cardiac dysfunction in pressure-overload model. **(A)** Ejection fraction (EF). 8-week-old mice were subjected to 27-gauge TAC or sham surgery and administered with vehicle (Veh), SLU-PP-332 (332) or SLU-PP-915 (915) by IP injection for 6 weeks. N=4-10, *: p<0.05 vs TAC+Veh group in each time point. **(B)** Gene expression of *Nppa* and *Nppb* in mouse heart tissue harvested after 6 weeks’ experiment. Expression levels were quantified by qPCR and normalized to Sham+Veh group. N=4-10, *: p<0.05 and ^#^: p<0.05. **(C)** Measurement of cardiac fibrosis. **Left panel**, representative fibrosis staining images using Masson Trichrome stain. Scale bar indicates 200 μm. **Right panel**, quantification of fibrotic area. N=4-10, *: p<0.05 and ^#^: p<0.05. **(D)** Representative transmission electron microscopy images of heart tissue section obtained from apex region after 6 weeks’ experiment. Arrow indicated mitochondrial fragmentation in TAC+Vehicle group. Scale bar indicates 500 nm. **(E)** Quantification of mitochondrial gene copy number upon 332 treatment. **Left panel**, copy number of *mt-Nd2* from mouse heart samples. N=4-6. *Chr6* was used as internal control. **Right panel**, copy number of *mt-Nd2* from NRVMs. N=3. *Chr4* was used as internal control. Statistical analysis was performed using two-way ANOVA for data in (A), and one-way ANOVA for data in (B), (C) and (E). Multiple comparison is corrected by Dunnet method for one-way ANOVA and Tukey method for two-way ANOVA, with α=0.05. Data are presented as mean ± SEM.

**Figure 2.**
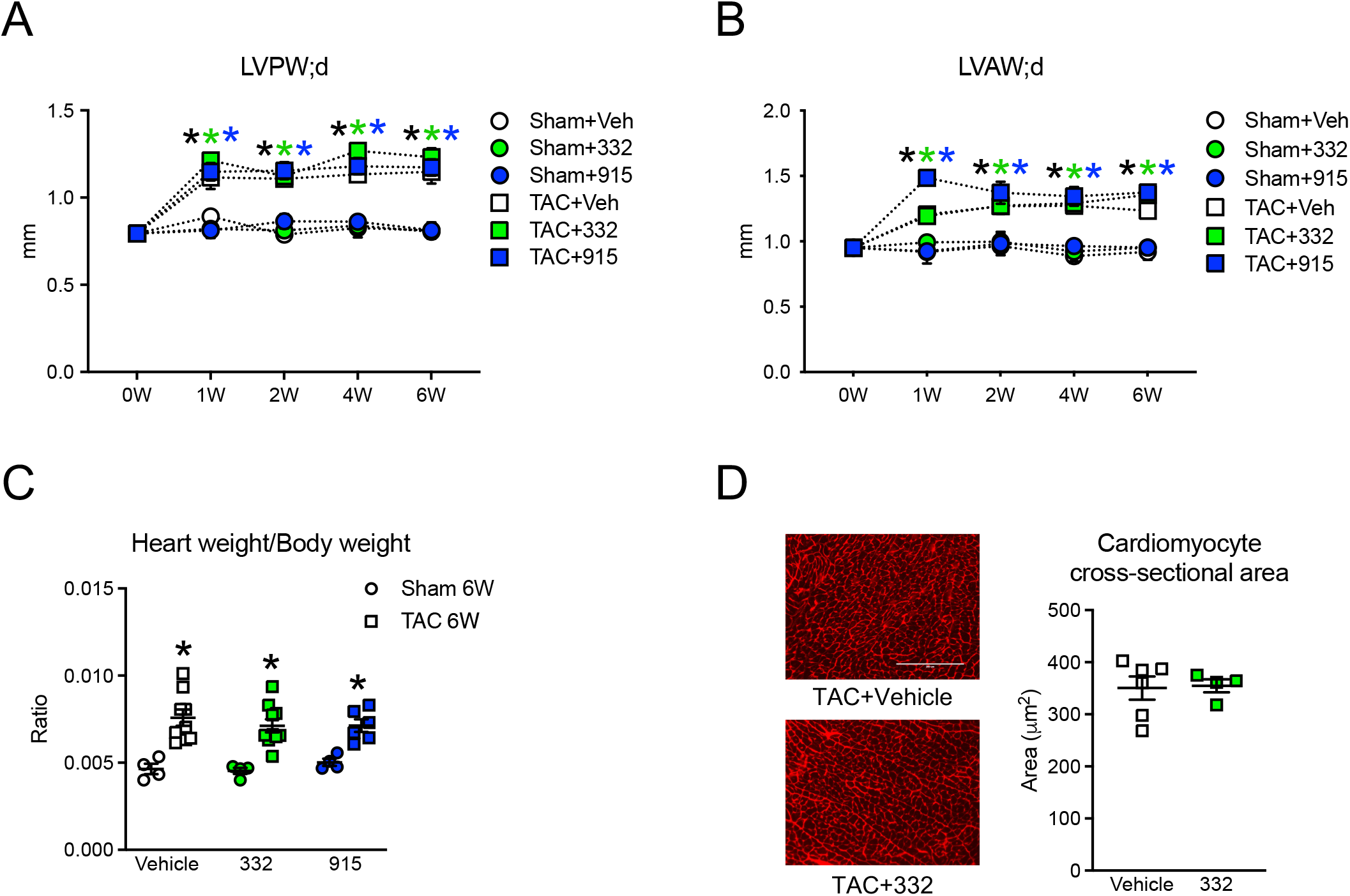
ERR agonists did not affect cardiac hypertrophy *in vivo*. **(A-B)** Left ventricular end-diastolic posterior wall thickness (LVPW;d) and left ventricular end-diastolic anterior wall thickness (LVAW;d). N=4-10, *: p<0.05 vs Sham+Veh group in each time point. **(C)** Ratio of heart weight to body weight. N=4-10, *p<0.05 vs Sham vehicle in each group. **(D)** Cardiomyocyte cross-sectional area measured by wheat germ agglutinin (WGA) staining. **Left panel**, representative images of cross-section with WGA staining for samples obtained after 6-week TAC. Scale bar indicates 200 μm. **Right panel**, quantification of cardiomyocyte cross-sectional area from the fluorescent images using imageJ. N=4-6. Statistical difference is determined by two-way ANOVA for data in (A), (B) and (C). Multiple comparison is corrected by Tukey method with α=0.05. Statistical analysis was performed using 2-tailed Student’s *t* test for data in (D). Data are presented as mean ± SEM.

Mitochondrial damage and dysfunction (39) play an important role in the pathogenesis of heart failure. ERR is known to participate in mitochondrial oxidative phosphorylation (OXPHOS) regulation; we then assessed whether 332 treatment reduced the mitochondrial damage in the mouse heart after 6-week TAC. Myocardial ultrastructure by electron microscopy in the TAC group showed an overall mitochondrial disarray with evident mitochondrial fragmentation and cristae destruction (Figure 1D), whereas administration of 332 greatly normalized the mitochondrial ultrastructure. This suggests that 332 prevented the mitochondrial damage seen in the TAC group. We also looked at the mitochondrial biogenesis by measuring the copy number of mitochondrial genes in both mouse heart samples and NRVMs. The results showed a mild trend of increase in the mitochondrial gene copy number in mouse heart treated with 332, and no apparent changes in NRVMs (Figure 1E and Supplementary Figure 2). Therefore, under our experimental condition, we did not observe a significant activation of mitochondrial biogenesis by 332 both *in vitro* and *in vivo*.

To confirm the cardioprotective effect provided by 332 was indeed by directly targeting ERR, we attempted the TAC rescue experiment with another novel ERR pan-agonist, SLU-PP-915 (915). 915 was designed utilizing a structure-based design approach taking advantage of the ligand bound crystal structure of ERRγ with known acyl hydrazide agonist, GSK-4716 (PDB:2GPP) (21). 915 has a distinct chemical scaffold from 332 (see chemical structureof 915 in Method). In cell-based cotransfection assay utilizing full length ERRs and a ERRE enhancer driven luciferase reporter, 915 showed a more balanced activity for both ERRα and ERRγ (E.C50 about 450nM) but lower activity toward ERRβ (E.C50 around 1.8uM, Supplementary Figure 3A). 915 was selective for the ERRs as it did not alter the activity of other nuclear receptors or GPCR (Supplentary Figure 3B-D). Pharmacokinetics of 915 were analyzed in mouse, showing plasma exposure (Supplementary Figure 4). We did not observe an overt toxicity in mice dosed at 25mg/kg during the entire course of the 6-week experiment.

In fact, 915 showed an even more pronounced cardioprotective effect post TAC than 332, as 915 significantly improved EF starting from 2 weeks post TAC (Figure 1A). Together, these data provide strong evidence suggesting that ERR agonists improve cardiac function and ameliorate mitochondrial damage and cardiac fibrosis against pressure overload induced-heart failure *in vivo*.

### ERR agonists did not ameliorate cardiac hypertrophy in vivo

Pressure overload in murine models consistently induces cardiac hypertrophy prior to reduction in cardiac function. Initially, the hypertrophy was thought to be adaptive in the early stage, but then eventually decompensates and becomes maladaptive due to chronic hemodynamic overload (40). It has also been suggested that hypertrophy per se may be a pathogenic signal (41). We investigated the effect of ERR agonists on cardiac hypertrophy in our TAC model. Considerable increase in left ventricular end-diastolic posterior wall thickness (LVPW;d) and left ventricular end-diastolic anterior wall thickness (LVAW;d) were observed as early as 1 week after TAC and sustained throughout the rest of the experiment (Figure 2A and 2B). Cardiac hypertrophy was also indicated by a higher heart weight to body weight ratio at 6-week post TAC (Figure 2C). Surprisingly, in contrast to the dramatic improvement seen in the cardiac function, mice administered with both 332 and 915 had similar levels of cardiac hypertrophy compared to that of the TAC group (Figure 2A and 2B) as well as similar heart weight to body weight ratios (Figure 2C). At the cellular level, cardiac hypertrophy results from an increase in individual cardiomyocyte size. Consistent with the gross echocardiogram and heart weight result, the cross-section area of cardiomyocytes delineated by wheat germ agglutinin (WGA) staining showed a similar degree of hypertrophy after 6-week TAC either treated with vehicle or 332 (Figure 2D). All these data suggest that ERR agonists did not prevent cardiac hypertrophy in the TAC model despite the significant improvement in cardiac function.

### ERR agonism did not suppress PE-induced activation of ERK1/2 and NFAT signaling *in vitro*

To further validate the *in vivo* results, we examined the effects of ERR agonism on cardiac hypertrophy *in vitro* and used an α-agonist phenylepinephrine (PE) to induce cardiomyocyte hypertrophy in NRVMs (Figure 3A). 24 hr treatment with PE increased the cell size by 37.6% in the vehicle group. Although 915 modestly reduced the cell size at baseline, PE still induced significant increases in cell size by 39.8% and 49.8% in the presence of 332 and 915, which agrees with the *in vivo* data. We further interrogated ERK1/2 signaling and calcineurin/NFAT signaling, which are two important signaling pathways that regulate cardiomyocyte growth and cardiac hypertrophy (42–45). As expected, PE significantly increased the level of ERK1/2 phosphorylation (Figure 3B). Both 332 and 915 had a modest to moderate induction of ERK1/2 phosphorylation at baseline. PE further stimulated the ERK1/2 phosphorylation in the 332 group while 915 appears to have a moderate suppression on PE-induced ERK1/2 phosphorylation. To further validate this result, we checked the gene expressions of *Fos* and *Egr1*, two immediate-early genes whose transcriptions are induced by ERK signaling (46–48). PE increased the expression of both *Fos* and *Egr1* in 24 hr (Figure 3C and 3D). 332 had no effects on PE-induced *Fos* and *Egr1* expression, while 915 attenuated the PE-induced *Fos* and *Egr1* expression. Taking these results together, 915 moderately suppressed the ERK1/2-induced cardiac hypertrophic signaling while 332 had little effect. To study calcineurin/NFAT signaling, we transiently transfected a NFAT-responsive luciferase reporter into the NRVMs and measured the NFAT transcriptional activity using the luciferase assay. Similar to what has been demonstrated previously (36), 24 hr PE treatment significantly induced the luciferase activity, suggesting an activation of calcineurin/NFAT signaling. However, treatment with 332 or 915 did not inhibit the activation of NFAT signaling by PE (Figure 3E). PE even had a stronger induction of NFAT activation in the 332 group. Therefore, both ERR agonists (332 and 915) improve cardiac function against heart failure but does not ameliorate the hypertrophic remodeling of the cardiomyocytes, particularly NFAT signaling-mediated pathological cardiac hypertrophy.

**Figure 3.**
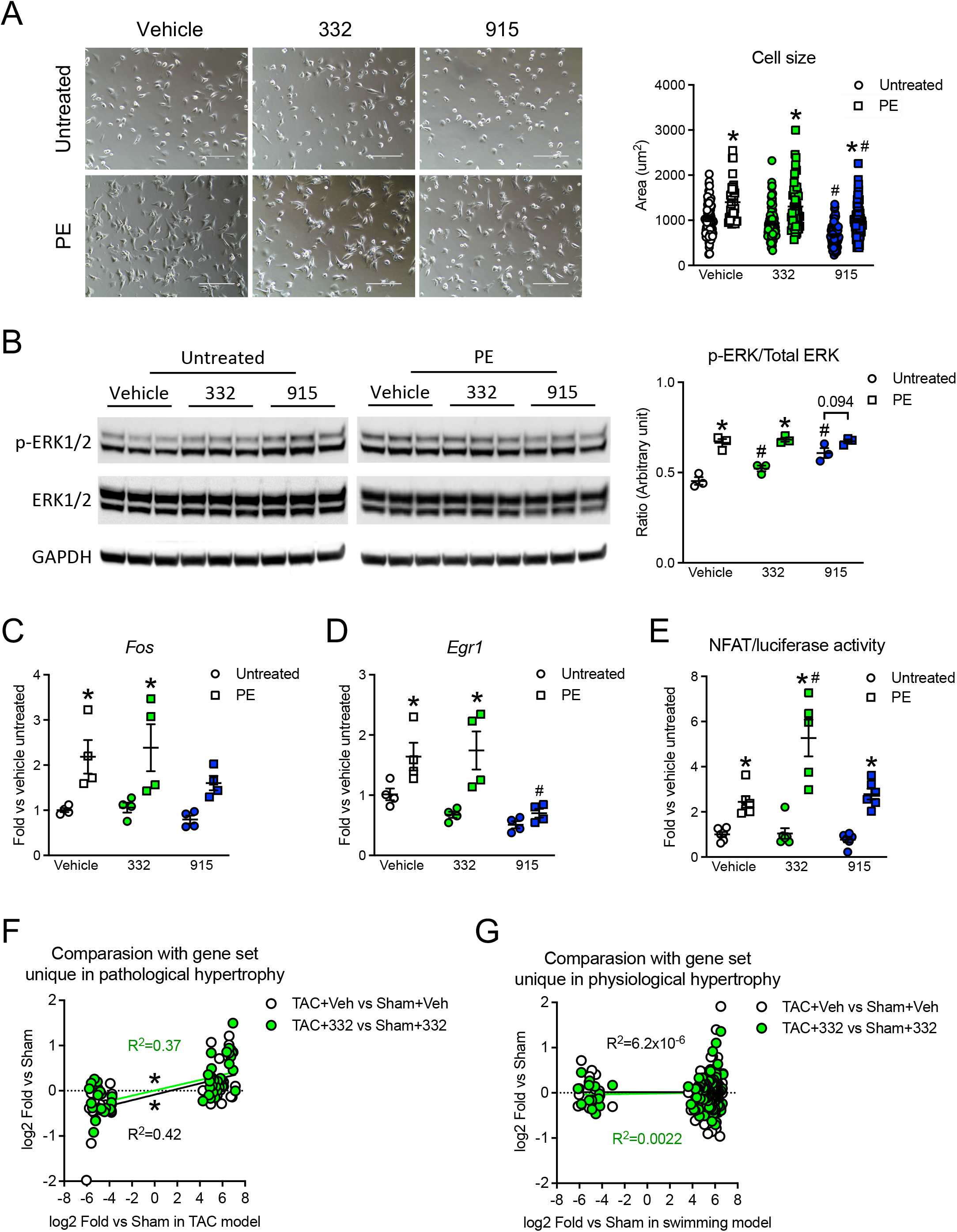
ERR agonism did not suppress PE-induced activation of ERK1/2 and NFAT signaling *in vitro*. **(A)** Cell size measurement in neonatal rat ventricular myocyte (NRVM). **Left panel**, representative images of NRVM treated with/without 100μM phenylephrine (PE) for 24hr in the presence of vehicle, 10μM 332 or 10μM 915. Scale bar indicates 200 μm. **Right panel**, quantification of cell size from bright field images using imageJ. N=18-51. **(B)** Western blot analysis of ERK1/2 signaling. **Left panel**, representative western blot images showing the protein levels of phosph-ERK1/2 (p-ERK1/2) and total ERK1/2 from NRVM. GAPDH were used as the loading control. **Right panel**, ratio of p-ERK1/2 to total ERK levels quantified from the western blot images. N=3. **(C)** and **(D)** Quantitative RT-PCR of ERK-targeted gene *Fos* and *Egr1*. Expression level of *Ppib* was used as internal control. N=4. **(E)** Dual luciferase reporter assay for NFAT transcriptional activity in NRVM. N=5-6. Statistic difference is determined by two-way ANOVA for data in (A), (B), (C), (D) and (E). *:p<0.05 and ^#^:p<0.05 vs untreated control within each group and untreated control in vehicle group, respectively. Multiple comparison is corrected by Tukey method with α=0.05. Data are presented as mean ± SEM. **(F)** and **(G)** Comparasion of gene sets specifically regulated in pathological or exersice model in mouse heart between a previous published study (49) and vehicle or 332 group. 137 and 52 genes were included in TAC-specific (F) and swimming-specific (G) gene sets, respectively. X-axis shows the fold changes in gene expressions between TAC and sham group (F) or swimming group and sham group (G) on a log-2 scale. Y-axis shows the fold change in gene expression between TAC and sham group with vehicle or 332 treatment on a log-2 scale, and the value represents average of three biological replicates. Linear lines across the data points indicate linear regression fitting. Asterisk indicates the slope of linear regression line is significantly different from zero.

### ERR agonism did not switch pathological hypertrophy to physiological hypertrophy in hearts with pressure overload

The fact that ERR agonism preserved the cardiac function but failed to prevent cardiac hypertrophy leads to test a hypothesis that ERR agonism may induce a transcriptional switch from pathological hypertrophy towards physiological hypertrophy despite of persistent pressure overload. We attempted to test this hypothesis by comparing our mouse heart RNA-seq data with two sets of genes that were previously reported to be unique to swimming-induced hypertrophy (physiological) or TAC-induced hypertrophy (pathological) from a qPCR-based screen in the mouse heart (49) (Figure 3F and 3G). We were able to match 137 swimming-specific genes and 52 TAC-specific genes to our mouse heart RNA-seq data set. To compare gene expression patterns, fold changes (vs sham) of each individual gene reported in the previous study were plotted along the x-axis on a log-2 scale, and fold changes (vs sham) calculated from our RNA-seq data were plotted along the y-axis on a log-2 scale. The results showed that expression profiles of both vehicle-TAC and 332-TAC groups correlate with TAC-specific genes expression profiles (Figure 3F), but not swimming-specific gene profiles previously reported (Figure 3G). Therefore, these data suggest that ERR agonism did not switch the pressure overload-induced pathological hypertrophy to physiological hypertrophy, instead they may have prevented the transition of compensatory hypertrophy to heart failure.

### RNA sequencing revealed ERR agonists upregulate fatty acid oxidation and OXPHOS metabolic pathways, and downregulate cell cycle and organ development pathways in cardiomycoytes

To further explore the molecular mechanism underlying the cardioprotective effects of ERR agonists in cardiomyocytes, which prevented the transition of compensatory hypertrophy to heart failure (50), we performed RNA sequencing first in NRVMs treated with 332 and 915. Overall, we identified 1137 (411 upregulated and 726 downregulated) and 2123 (602 upregulated and 1521 downregulated) differentially expressed genes (DEGs) in the 332 and 915 treated NRVMs respectively with a cutoff of 1.5-fold compared to the vehicle group (Figure 4A). 879 genes were found to be differentially expressed in both the 332 and 915 groups, where 244 genes were upregulated and 624 genes were downregulated. Significant overlap of DEGs between the 332 and 915 groups is observed as expected, confirming their shared ERR targets and high specificity.

**Figure 4.**
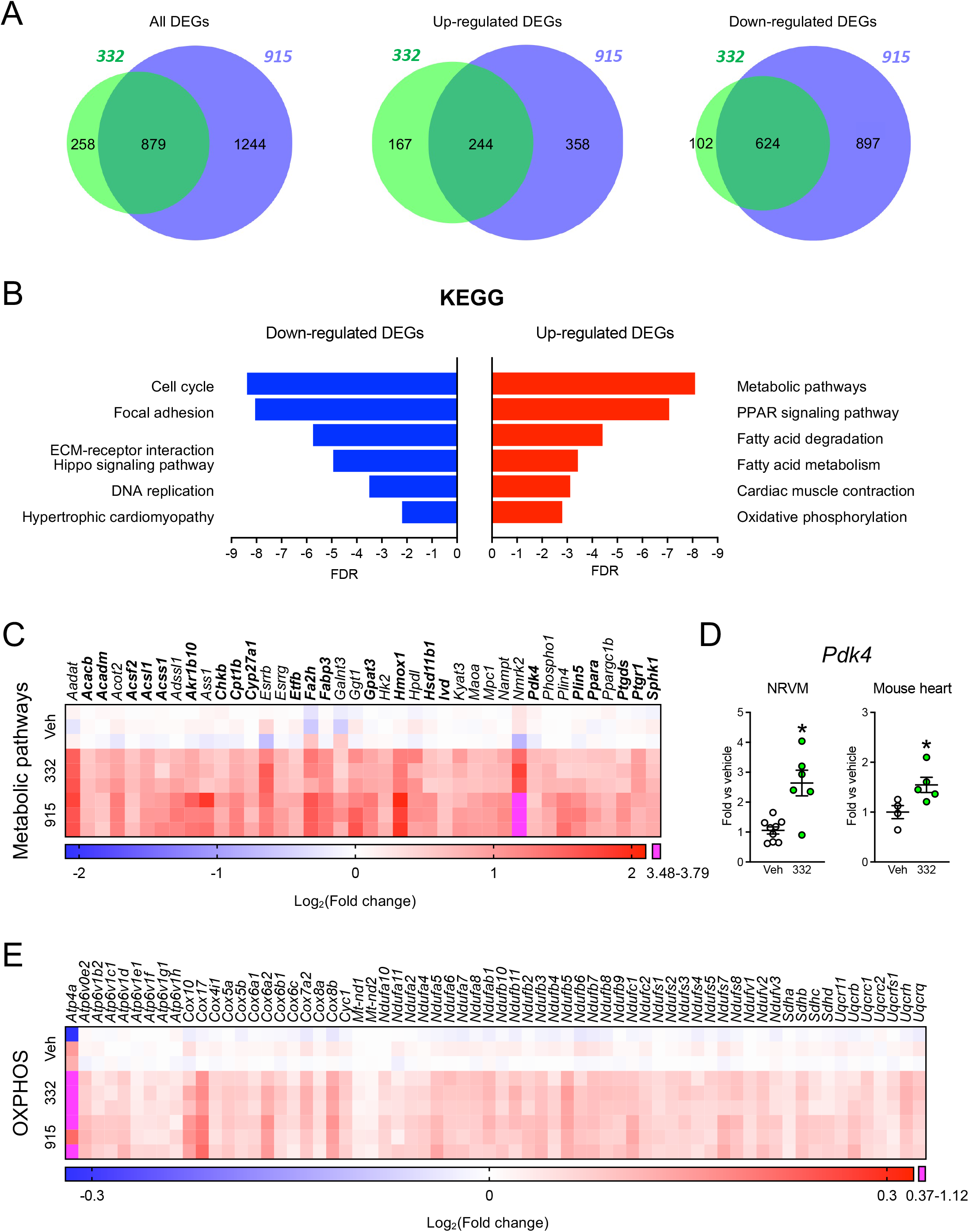
RNA sequencing revealed ERR agonists upregulate fatty acid metabolic pathways and downregulate cell cycle and organ development pathways in cardiomyocytes. RNA sequencing was performed on NRVMs treated with vehicle, 10μM 332, or 10μM 915 for 72hr. Differentially expressed genes (DEGs) with 1.5-fold difference versus vehicle group and adjusted p value<0.05 were included in the analysis. **(A)** Venn diagram comparing DEGs between NRVMs treated with 332 (green) and 915 group (purple). **(B)** KEGG pathway analysis of upregulated and downregulated DEGs pooled from the 332 and 915 groups. FDR (false discovery rate) was shown in x-axis. **(C)** Heatmap of metabolic-related DEGs commonly upregulated by both 332 and 915. Genes involved in fatty acid/lipid metabolism were highlighted in bold. Genes with 1.5-fold increase upon 332 or 915 treatment vs vehicle group and adjusted p value<0.05 were included in the analysis. N=3. Scale bar indicates a range of Log_2_(Fold change) from −2.1 to 2.1 (0.233-4.287 fold). Magenta indicate Log_2_(Fold change) values between 3.48-3.79 (11.158-13.833 fold). **(D)** 332-induced gene expression of *Pdk4 in vitro* and *in vivo*. Left panel, *Pdk4* expression level in NRVMs treated with vehicle or 10μM 332 for 72hr. Right panel, *Pdk4* expression level in mouse treated with vehicle or 25mg/kg 332 twice a day for 10 days. Expression level of *Ppib* was used as internal control. N=4-9. Statistical difference was determined by 2-tailed Student’s *t* test. Data are presented as mean ± SEM. **(E)** Heatmap of genes involved in oxidative phosphorylation (OXPHOS). Genes with significant changes in expression levels (adjusted p value<0.05) upon 332 or 915 treatment vs vehicle group were included in the analysis. N=3. Scale bar indicates a range of Log_2_(Fold change) from −0.32 to 0.32 (0.801-1.248 fold). Magenta indicate Log_2_(Fold change) values between 0.37 to 1.12 (1.292-2.173 fold).

To further interrogate the RNA-seq data, we first performed pathway enrichment analysis on the DEGs in the 332 and 915 groups separately. We found that DEGs in the 332 and 915 groups showed similar enrichments from gene ontology analysis and KEGG pathway analysis (Supplementary Figure 5), indicating that 332 and 915 indeed regulate similar biological pathways. Therefore, we pooled the DEGs in both the 332 and 915 groups together and analyzed the pooled DEGs, which consists of 769 upregulated genes and 1623 downregulated genes. KEGG Pathway analysis showed that upregulated genes are enriched in metabolic pathways, particularly fatty acid/lipid metabolism (Figure 4B). Not too surprisingly, we also observed an enrichment in PPAR (peroxisome proliferator-activated receptor) signaling pathways, as both ERR and PPAR regulate fatty acid metabolism (16). Significant enrichment in cardiac muscle contraction and oxidative phosphorylation (OXPHOS) were also observed in upregulated DEGs. ERRs are known for their transcriptional activation activity in fatty acid oxidation (FAO) and mitochondrial genes. We were surprised to see an even larger number of downregulated DEGs. Downregulated DEGs are markedly enriched in pathways that are essential for cell proliferation and developmental processes, such as the cell cycle, focal adhesion, ECM-receptor interaction, and DNA replication. Notably, enrichment in Hippo signaling pathways, which is vital for organ size development (51), was also observed. Mature cardiomyocytes are terminally differentiated cells and have very limited ability to re-enter the cell cycle (52). Reactivation of cell cycle and fetal gene expression has been causally linked to the development of cardiomyopathy and heart failure (53–55); thus, suppression of fetal gene expression could be beneficial for heart failure. We also noticed that downregulated DEGs were enriched for hypertrophic cardiomyopathy; however, we did not observe significant effects of ERR agonism on cardiac hypertrophy (Figure 2 and Figure 3). Our data is largely in agreement with early microarray studies in the ERRα and ERRγ knockout mouse models (9, 11, 12).

A previous study using ChIP-on-chip analysis identified 195 and 231 promoters bound by ERRα and ERRγ in the mouse heart (11). When compared with the list of DEGs from our RNA-seq data, we only found 32 genes that overlap with DEGs in the 332 or 915 groups (Table 1). The small number of overlapping genes suggests that a large fraction of the DEGs are indirect targets of ERR, or the regulatory sequences are located in enhancers far away from the regulated targets. This may also be attributed to differences between rat and mouse species as well as to the relatively low sensitivity of the ChIP-on-chip technique. Nonetheless, it is interesting that among these 32 genes (20 upregulated, 12 downregulated), most of the upregulated genes are metabolic genes while the downregulated ones are non-metabolic genes. These data agree with previous studies showing that metabolic genes are the primary targets of ERR (9–13), meanwhile, our data also raise the possibility that ERR may repress directly or indirectly transcription of non-metabolic genes.

To further validate our findings, we performed RNA-seq study on mouse heart tissue to examine the perturbation of cardiomyocytes upon ERR agonism *in vivo*. Because heart tissue consists of both cardiomyocyte and non-cardiomyocyte (fibrobaslat, endothelial cells, immune cells), we applied deconvolution of bulk-RNA seq profile of whole heart tissue to isolate the expression profile of each invidual cell types using XDec (Supplementary Figure 6). XDec deconvolution is a reference-free deconvolution method that allows us to discern constituent cell types, their proportions, and cell type-specific expression profiles in heterogenous biological samples (56). The method also has the potential to identify multiple distinct states of the same cell type, such as myocyte, pre- and post-treatment.

Deconvolution of heart bulk-RNAseq profiles of vehicle/332 treated sham/TAC mouse heart samples identified four cardiomyocyte cell states with unique expression profiles (Profile 1-4 in Supplementary Figure 6A). Meanwhile, only one of each profile was identified for immune cell-like profiles (Profile 7), endothelial-like (Profile 8), and cardio-fibroblast-like (Profile 9), which may indicate that ERR agonism primarily targets the cardiomyocyte. Profile 3 and 4 were found to be enriched in vehicle control (83.06%) and 332 treated (69.68%) TAC samples, respectively (Supplementary Figure 6B; Two-sample t-test, p=0.047 and p=0.024, Supplementary Figure 6C). Therefore, profile 3 and 4 likely represent the cardiomyocyte populations in vehicle control and 332 treated TAC hearts, respectively. To determine the effect of ERR agonism on gene expression of cardiomyocytes, we then compared the average expression profiles of these two constituent cell types and identified 1553 upregulated and 762 downregulated DEGs. KEGG pathway analysis of those DEGs showed enrichment of OXPHOS genes in up-regulated DEGs upon 332 treatment (Supplementary Figure 6D). Genes associated with adrenergic signaling and cardiac muscle contraction were also enriched in the up-regulated DEGs. Among downregulated DEGs, we found significant enrichment in the Hippo signaling pathway, which is consistent with our finding *in vitro* (Figure 4B and Supplementary 5B). Overall, these results aggree with the RNA-seq results from NRVMs, and corroborate with our *in vitro* findings that ERR agonist elevates the metabolic funcntion in cardiomycoytes. On the other hand, ERR agonism represses HF-induced fetal gene expression, in which Hippo signaling pathway may play an important role.

### ERR agonism induces transcription of FAO and OXPHOS-related genes and increases oxidative metabolism in cardiomyocytes

Since pathway analysis indicates ERR agonism activates metabolic pathways in cardiomyocytes, we further analyzed the DEGs that are upregulated by both 332 and 915 *in vitro*. From the pathway analysis, we identified 40 metabolic related genes that are commonly targeted by both 332 and 915 *in vitro* (Figure 4C). In line with the pathway enrichment analysis, 22 of the 40 genes participate in fatty acid/lipid metabolism (Bold in Figure 4C), including both specific enzymes such as *Acacb*, *Acadm*, *Acsl1*, and *Cpt1b* and major metabolic regulators, such as *Pdk4*, *Ppara*, and *Ppargc1b.* In particular, *Pdk4* is a known ERR target gene which plays a pivotal role in controlling the metabolic plasticity (57–59). Increased transcription of *Pdk4* leads to the energy source switching from glucose to fatty acid in the heart and other metabolically active tissues (60, 61). We validated the increased transcription of *Pdk4* in NRVMs and mouse heart upon ERR agonism using 332 (Figure 4D). Therefore, PDK4 is likely an essential mediator for boosting fatty acid metabolism upon ERR agonism. Moreover, treatment of 332 or 915 led to a moderate increase in expression level of OXPHOS genes (Figure 4E), which is also consistent with the pathway analysis. In addition, we also observed moderate increases in genes involved in the TCA cycle and glycolysis (Figure 4C and Supplementary Figure 7).

To directly examine the metabolic effects of ERR agonists at the functional level, we analyzed the metabolomics profile of mouse hearts that underwent 6-week TAC with or without 915 treatment. The results showed that 6-week TAC led to a pronounced alteration in most of the metabolite levels that we examined. 915 treatment significantly attenuated the TAC-induced perturbation in metabolite levels (Figure 5A). Remarkably, 112 lipid/fatty acid metabolites were decreased after 6-week TAC, while only 5 were found decreased when treated with 915. This result agrees with the RNA-seq data and further emphasizes that fatty acid metabolic pathways are a primary target of ERR. In addition, 46.2% of TCA/OXPHOS metabolites (6 out of 13) were altered after 6-week TAC, but only 7.7% (1 out of 13) were changed in the 915 group, indicating a normalization of TCA/OXPHOS metabolic pathways by ERR agonism. The normalization of metabolite levels was also observed in other metabolic categories that we examined, including carbohydrate, amino acid, nucleotide, peptide, cofactor and xenobiotics, although TAC induced changes in these pathways were more limited. Therefore, both transcriptomic and metabolomic data strongly suggest that ERRs agonists provide cardioprotection through benefical cardiac metabolic remodeling, particularly by boosting fatty acid metabolism and OXPHOS.

**Figure 5.**
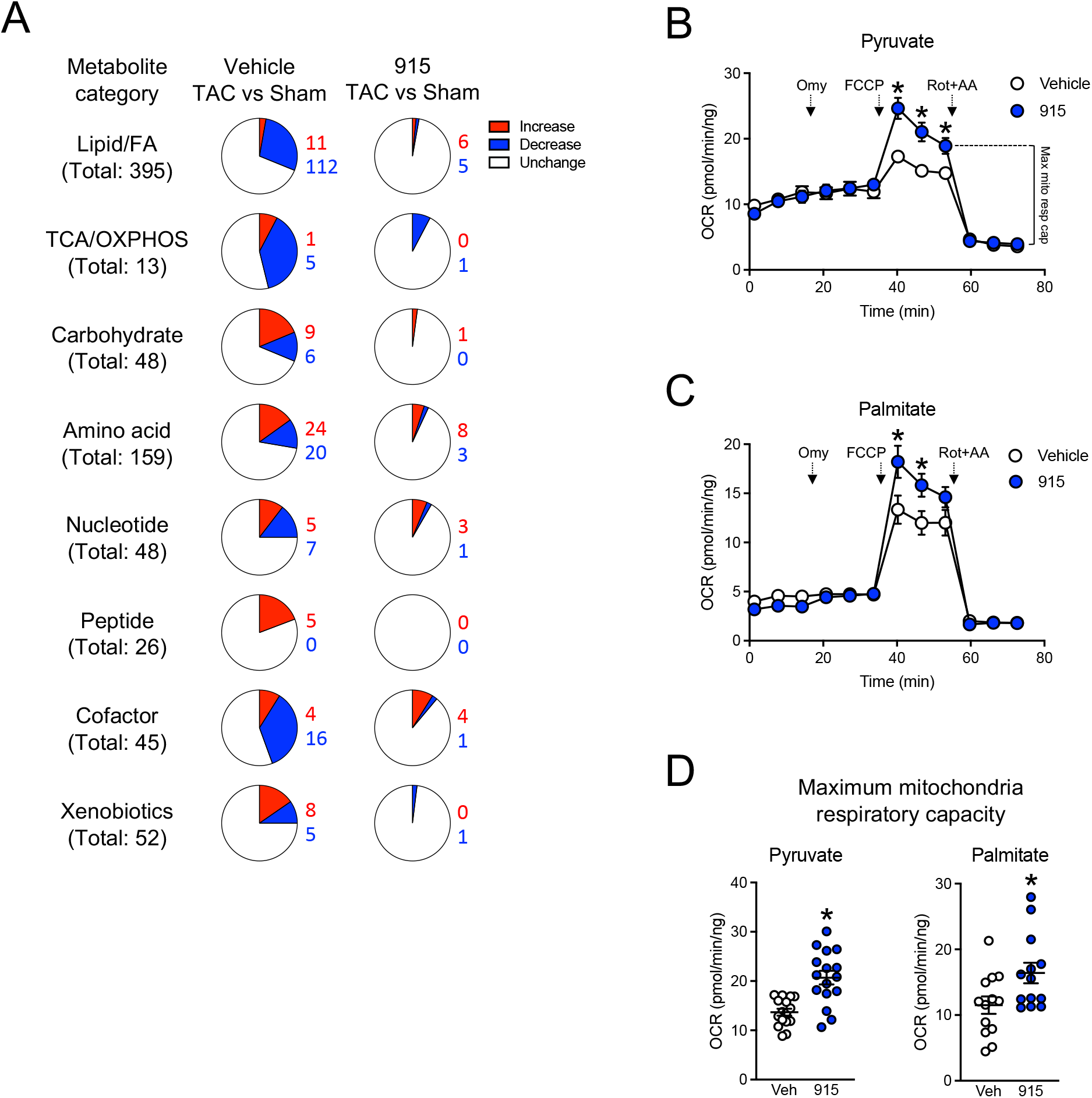
ERR agonism increased the oxidative metabolism in cardiomyocytes. **(A)** Metabolomics analysis obtained from mouse heart tissue after a 6-week TAC experiment. Numbers in blue or red next to the pie chart indicate the number of metabolites that are decreased or increased compared to the Sham group. N=4-8. **(B-C)** Measurement of oxygen consumption rate of isolated adult mouse cardiomyocytes in extracellular flux analyzer XFe96. Pyruvate (B) and palmitate (C) were used as substrate to assess the OXPHOS function. Arrows indicate where drugs were injected. Error bars are not shown when smaller than the symbol size. **(D)** Quantification of maximum mitochondria respiratory capacity. **Left panel**, data obtained with pyruvate as the substrate. N=16. **Right panel**, data obtained with palmitate as the substrate. N=13. *:p<0.05. Statistical differences between 915 and vehicle group were determined by two-way ANOVA for data in (B) and (C), and 2-tailed Student’s *t* test for data in (D). Multiple comparison is corrected by Tukey method with α=0.05. Data are presented as mean ± SEM.

Finally, to assess the metabolic flux of fatty acid and OXPHOS metabolic pathways, we used palmitate or pyruvate as the substrate and measured oxygen consumption rate (OCR) of cardiomyocytes isolated from adult mouse hearts (ACMs) (Figure 5B-D). We found that ACMs treated with 915 showed significantly elevated maximum mitochondria respiratory capacity with either pyruvate or palmitate as a substrate, which further provides functional confirmation that ERR agonists upregulate the activity of oxidative metabolism capacity in cardiomyocytes. In conclusion, ERR agonists may improve overall metabolic gene profiling and maintain cellular energy level in cardiomyocytes with the most prounced modulation on activation of fatty acid metabolism.

### ERR agonism induced autophagy in cardiomyocyte

Activation of autophagy is generally considered to be cardioprotective in a variety of cardiovascular disease settings (62). We found that ERR agonism induced a mild upregulation of autophagy-related genes at the transcript level (Figure 6A). Consistently, 72hr treatment of 332 or 915 led to increased protein levels of LC3-II or p62 in NRVMs at baseline condition (Figure 7B). We further treated NRVMs with bafilomycin which inhibits autophagosome-lysosome fusion to assess autophagic flux (63). The results showed that 915 elevated both LC3-II and p62 levels, while 332 increased p62 but not LC3-II level (Figure 7B), indicatin 915 may be a more potent autophagy activator than 332. Collectively, these data suggest that in addition to improving the metabolic function, ERR agonism also induced autophagy in cardiomyocyte.

**Figure 6.**
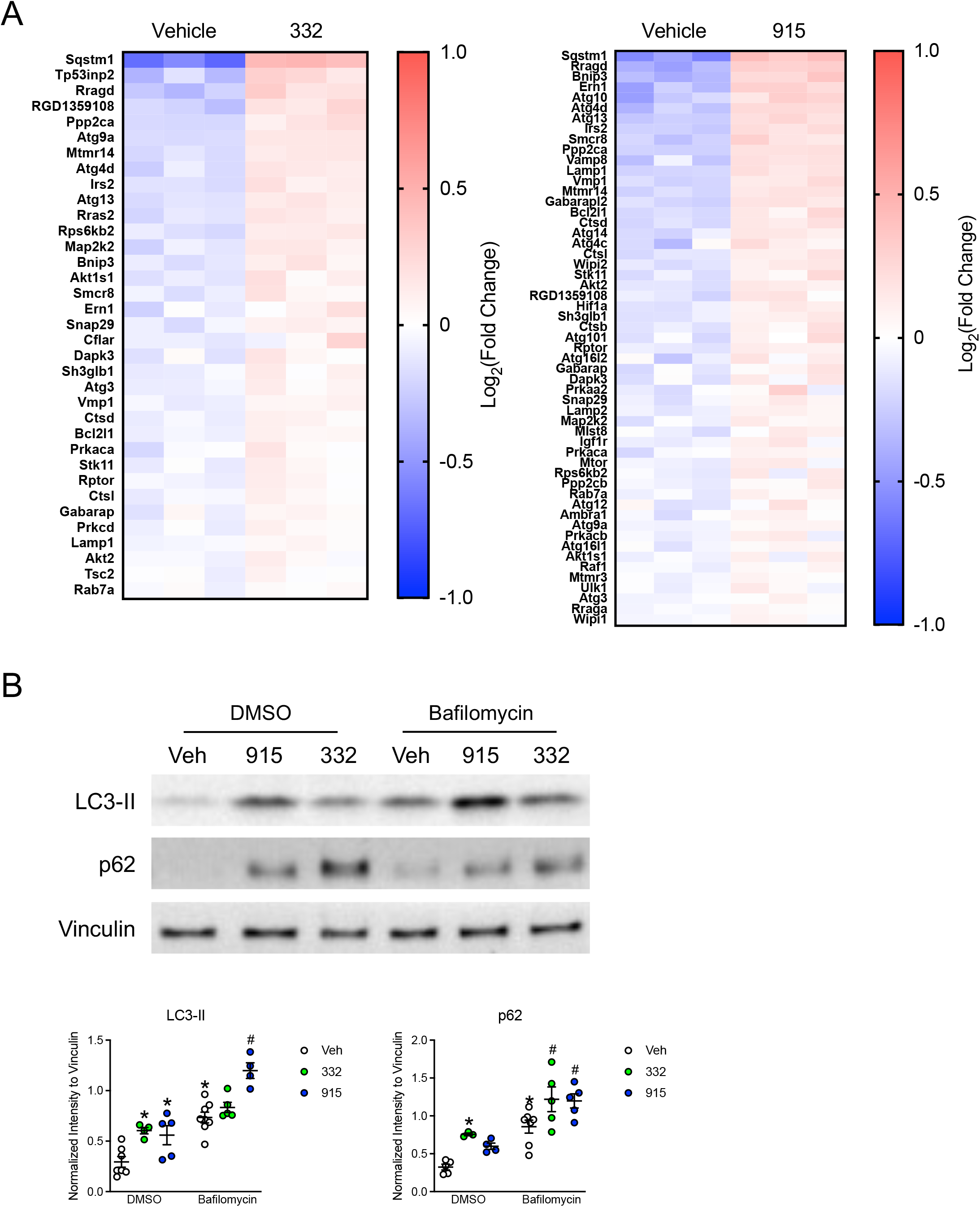
ERR agonism leads to activation of autophagy in cardiomyocyte. **(A)** Heatmap of genes involved in autophagy pathway and upregulated by 332 and 915 from NRVM RNAseq. Only genes with significant changes were included (adjust p-value<0.05). N=3. **(B)** Western blot analysis of LC3B and p62 in NRVM treated with vehicle, 5μM 332 or 2.5μM 915 for 72hr. Cells were treated with additional 2hr of 100nM bafilomycin (Cell Signaling Technologies, 54645) to assess autophagic flux. The concentration of 332 or 915 was titrated to minimize cell death in the assay. **Upper panel**, representative of western blot images. **Lower panel**, quantitative of LC3B or p62 protein level from the western blot images. Vinculin were used as loading control. N=3-8. *p<0.05 vs vehicle in the control group. ^#^p<0.05 vs vehicle in the bafilomycin group. Statistical difference is determined by two-way ANOVA. Multiple comparison is corrected by Tukey method with α=0.05. Data are presented as mean ± SEM.

**Figure 7.**
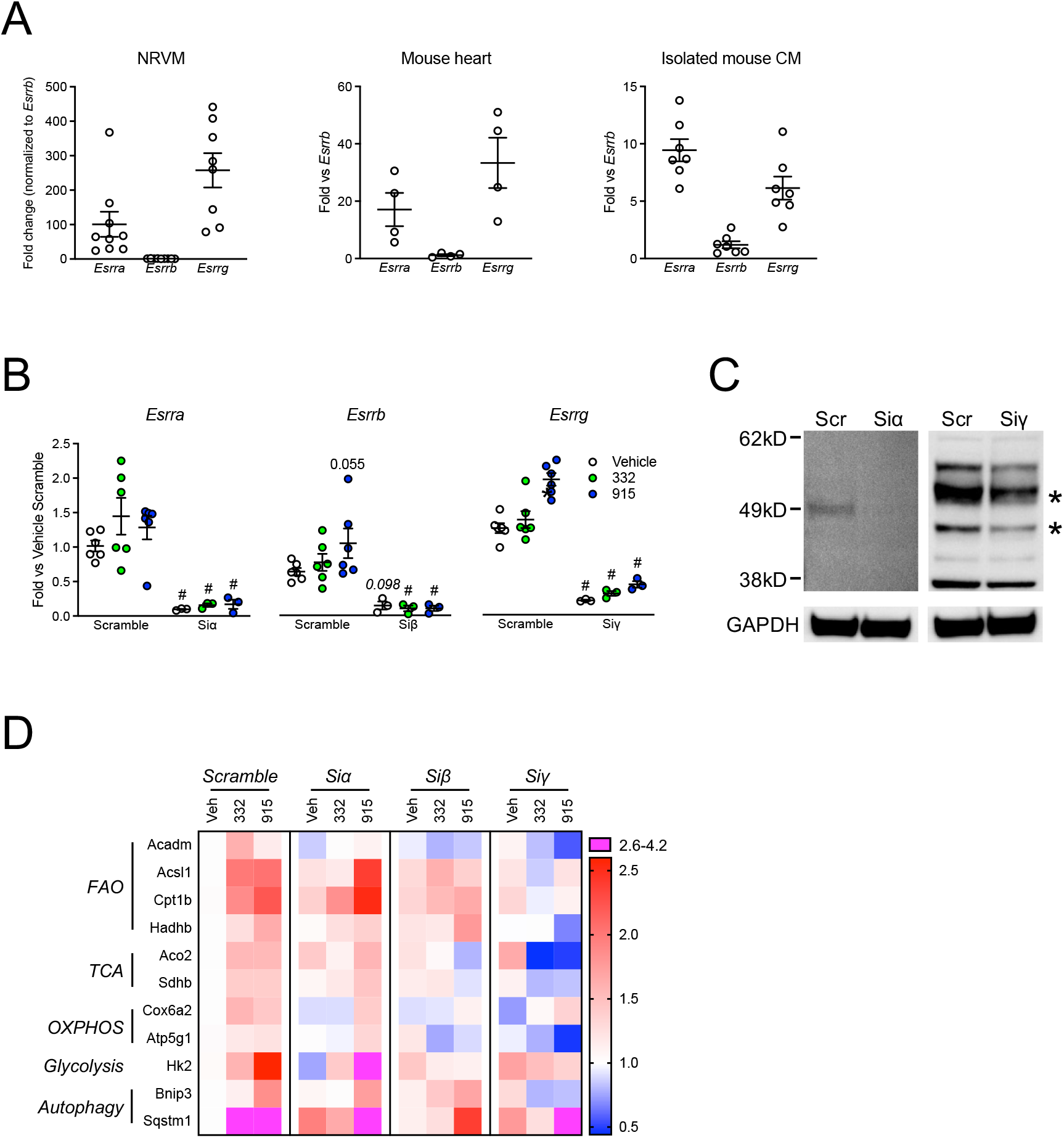
ERRγ is the major mediator for transcriptional regulation by ERR agonists in cardiomyocytes. **(A)** Gene expression level of three ERR isoforms in NRVMs, mouse heart and isolated mouse cardiomyocytes. Gene expression level was quantified by qPCR and normalized to *Esrrb*. N=4-9. **(B)** Gene expression of three ERR isoforms upon siRNA silencing. NRVMs were harvest 72hr after transfected with 25pM scramble or ERR siRNA. Gene expression level was quantified by qPCR and normalized to the vehicle scramble group. N=3-6. **(C)** Western blot analysis of ERRα and ERRγ expression in NRVM 72hr after transfected with scramble or ERR siRNA. Representative western blot images were shown to demonstrate knockdown of ERRα and ERRγ at protein level. Asterisks indicates the bands of two ERRγ isoforms. The other bands were non-specific. GAPDH were shown as loading control. **(D)** Heatmap summarizing the effects of ERR isoform knockdown on ERR agonist-induced gene expressions. NRVMs were transfected with 25pM scramble or ERR siRNA. 24hr after transfection, NRVMs were treated with vehicle, 10μM 332, or 10μM 915 for 24hr. Gene expression level was quantified by qPCR and normalized to the vehicle scramble group. N=3-6. Expression level with a fold change of 2.6 to 4.2 compared vehicle scramble group is indicated in pink. Statistical difference is determined by one-way ANOVA for data in (A) and two-way ANOVA for data in (B). Multiple comparison is corrected by Dunnet method for one-way ANOVA and Tukey method for two-way ANOVA, with α=0.05. Data are presented as mean ± SEM.

### ERR agonist-induced transcriptional regulation is mainly mediated by ERRγ

All three ERR isoforms are expressed in NRVMs, mouse hearts, and isolated mouse cardiomyocytes, with ERRα and ERRγ being the dominant isoforms (Figure 7A). Both 332 and 915 are ERRs pan-agonists. To further dissect the main ERR isoform mediating the cardiac protective effects, we performed siRNA knockdowns for each ERR isoform individually to interrogate ERR dependency for 332 and 915-induced effects on gene transcription (Figure 7B). Efficient knockdown was achieved by reducing transcript level by 90.4%, 77.2%, and 83.3% for *Esrra*, *Esrrb,* and *Esrrg* respectively at baseline conditions (Figure 7B). In the control (scrambled) group, we found that 915 increased expression levels of *Esrrb* and *Esrrg* (Figure 7B), which is consistent with the RNA-seq data (Figure 4C) and further suggests a positive feedback mechanism may be activated upon ERR stimulation. Knocking down of ERRα and ERRγ was further confirmed at the protein level by immunoblotting (Figure 7C). We failed to detect an ERRβ band using commercially available antibodies in the NRVMs, which is likely due to its low expression level in the cardiomyocytes (Figure 7A).

We then selected 11 key metabolic genes with considerable alterations in expression level based on the RNA results for investigation, including *Acsl1*, *Cpt1b*, *Acadm* and *Hadhb* for fatty acid metabolism, *Sdhb* and *Aco2* for the TAC cycle, *Hk2* for glycolysis, *Cox6a2* and *Atp5g1* for oxidative phosphorylation (Figure 7D and Supplementary Figure 8A). Notably, *Hk2* and *Acadm* have been suggested as direct targets of ERRα and ERRα/γ (Table 1) (11). Two important autophagy genes that are predicted to have considerable changes in transcription level by RNA-seq data, *Sqstm1* and *Bnip3,* were also included (Figure 7D and Supplementary Figure 8B). As a validation of the RNA-seq result, all the genes showed significantly increased expression levels upon 332 or 915 treatment in the scramble group, except for *Atp5g1,* which also showed a trend of increase. We noticed that while treatment with 332 and 915 generally exhibited the same trend, their individual effect on particular genes varies, e.g. *Acadm*, *Hadhb*and *Hk2*, which may reflect subtle differences in potency or isoform specificity of these two compounds.

Next, three ERR isoforms were individually knocked down to examine the ERR dependency of ERR agonist-induced transcription in the 10 genes shown in Figure 7D (Also in Supplementary Figure 8). The results demonstrated that knocking down of ERRγ significantly prevented 332 or 915-induced gene expression in all the genes except 915-induced *Cox6a2* expression. To our surprise, knocking down of ERRα only diminished 332 or 915-induced gene expressions in *Acsl1*, *Acadm,* and *Cox6a2* but had no apparent effects in the other genes tested. Dysfunction of ERRβ has been associated with pathogenesis of dilated cardiomyopathy by affecting contractility and calcium balance (64), however, whether ERRβ regulates cardiac metabolic function has not been explored yet. Interestingly, despite of its low expression level in NRVMs, we observed ERRβ-depdendent transcriptional activation by 332 or 915 in *Acsl1*, *Cpt1b*, *Sdhb*, *Acadm*, *HK2*, *Aco2*, *Cox6a2,* and *Sqstm1*. In addition, although 332 and 915 only induced a trend of increase in *Atp5g1* expression, we found its expression level appears to be lower when ERRβ or ERRγ was knocked down. To facilitate visualization of the result, we summarize the ERR isoform dependency qRT-PCR results in a heatmap, which shows the expression pattern was most significantly altered in the ERRγ knockdown group (Figure 7D). Therefore, all three ERR isoforms participate in the transcriptional regulation by ERR agonism, where ERRγ appeared to be the main mediator at least in the panel of selected genes tested. Our data further revealed that despite the low expression levels, ERRβ may also play important roles in regulating cardiac metabolism and may be a novel therapeutic target for heart failure treatment (15, 16). Future work combining pharmacological agonists with isoform specific mutant mice will be informative.

### Downregulation of cell cycle genes by ERR agonism is partially mediated by E2F1

To address whether suppression of cell cycle genes is due to direct transcriptional regulation through ERR, or a consequence of altered cardiac metabolism, we did a motif analysis for the promoter regions (-1000 to 0 bp) of the downregulated DEGs. The results showed that the top enriched motif is E2F transcription factor binding site in both the 332 and 915 groups (Figure 8A). E2F or the adenoviral early region 2 binding factor is a major transcriptional regulator for cell cycle and DNA synethesis in mammalian cells. The E2F family consists of 9 member, in which E2F1/2/3A are transcriptional activators and E2F3B/4/5/6/7/8 are transcriptional suppressors in mammalian cells. Activation of different E2Fs requires binding with distinct cofactors such as transcription factor dimerization partner family for E2F1-6, thereby confers distinct mechanism of action (65). We set out to search for potential ERR binding sites in the promoter regions (-1000 to 0 bp) of the eight *E2F* members, and found three ERRα binding sites in *E2f1* (-16, -45, and -631 bp), two binding sites (-63 and -218 bp) in *E2f2*, one binding site (-241 bp) in *E2f3*, but none near any of the suppressors. In addition, the RNA-seq data demonstrated that the gene expression level of *E2f1*, but not *E2f2* or *E2f3*, were significantly reduced in NRVMs after treated with 332 or 915. Therefore, *E2f1* is the top candidate that may mediate the downregulation of cell cycle genes upon ERR agonism.

**Figure 8.**
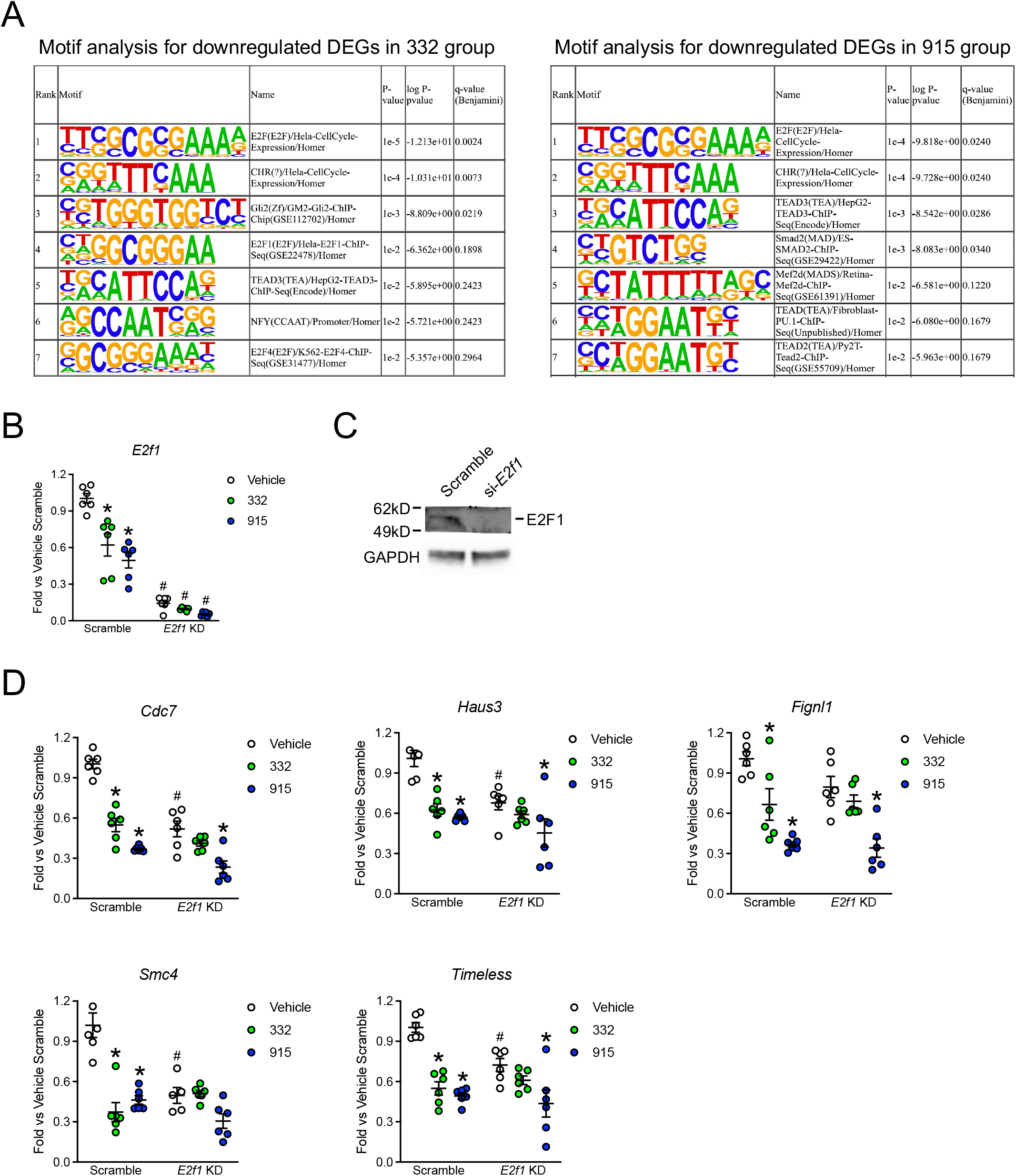
E2F1 is an important mediator for the downregulated DEGs upon ERR agonism *in vitro*. **(A)** Motif analysis for downregulated DEGs in both the 332 and 915 group from NRVM RNAseq. Top seven enriched motifs were shown in the tables. **(B)** Gene expression of *E2f1* upon *E2f1* siRNA silencing. N=6. **(C)** Western blot analysis of E2F1 expression in NRVM 72hr after transfected with scramble or *E2f1* siRNA. Representative western blot images were shown to demonstrate considerate knockdown of E2F1 at protein level. **(D)** E2F1-dependent DEGs. N=6. Downregulated DEGs with predicted E2F1 cognate site and over 1.5-fold change in expression upon ERR agonism were selected and examined in the experiment. Statistical differences were determined by two-way ANOVA for data in (C) and (C). Multiple comparison is corrected by Tukey method with α=0.05. Data are presented as mean ± SEM.

To further deciper the role of E2F1 in regulating the cell cycle genes in the cardiomyocytes, we performed knockdown of *E2f1* expression by siRNA in NRVMs. Efficienct knockdown was achieved by 86% reduction of the expression at baseline condition (Figure 8B). Knocking down of E2F1 was further confirmed at the protein level by immunoblotting (Figure 8C). Concordant with the RNA-seq data, treatment of 332 and 915 largely decreased the expression level of *E2f1* (Figure 8B). Based on the motif analysis and the RNA-seq data, we selected 15 cell cycle genes which contain E2F1 cognition sites in the promoter regions and exhibited significant downregulation in both the 332 and 915 groups in the RNA-seq data for further investigation. Quantitative PCR analysis for the control group (scramble) validated the suppression of transcription in all the 15 genes after ERR agonism (Figure 8D and Supplementary Figure 9). With *E2f1* knockdown, all the genes showed decreased expression levels compared with that in the scramble groups, indicating that these genes are indeed transactivated by E2F1 at baseline condition. The 332-induced downregulations of gene transcription were diminished in *Cdc7*, *Fignl1*, *Haus3*, *Smc4*, and *Timeless* with *E2f1* knockdown (Figure 8D). *E2f1* knockdown also attenuated 915-induced downregulation of *Smc4* transcription (Figure 8D). The reduced expression of the other 10 genes after ERR agonism was not affected, suggesting the ERR agonists induced reduction in these genes are through E2F1-independent mechanisms (Supplementary Figure 9). Therefore, E2F1 partially mediates the downregulation of cell cycle gens upon ERR agonism, additional mechanisms also exist.

## Discussion

We showed 2 novel structurally distinct ERR pan-agonists, 332 and 915, protected cardiac function against pressure overload-induced heart failure in a murine model, which is a critical step towards developing ERR agonists for heart failure therapy. The highly overlapping DEGs post 332 and 915 treatment is consistent with high selectivity in these two compounds (Figure 4A and Supplementary Figure 3).

The mammalian heart expresses three ERR isoforms with ERRα and ERRγ expressing at much higher levels than ERRβ (Figure 7A) (10). The essential role of ERRα and ERRγ in controlling cardiac metabolism, mitochondrial and contractile function has been demonstrated by early studies of the genetic knockout mouse model (9–13). ERRα and ERRγ were shown to bind to similar promoters and share a significant proportion of their target genes (11). Cardiac-specific knockout of ERRβ in mice lead to dilated cardiomyopathy with impaired calcium handling at 9 months of age, suggesting that ERRβ also plays a role in cardiac contractile function (64). However, the exact molecular mechanism and direct targets of ERRβ were not reported to date to our knowledge. While ERR-targeting therapy has been proposed for heart failure therapy, no natural ligand was identified nor synthetic agonist with favorable pharmacokinetics that can be used for *in vivo* studies had been developed (15, 16). Although there have been compounds identified as ERRα agonists (17, 18), ERRγ agonists (19), and ERRβ/γ agonists (20) through chemical library screening, none of them have been available for *in vivo* use and examined as a therapeutic agent for heart failure treatment *in vivo*. We believe that it was necessary to develop a chemical “tool” compound that would have *in vivo* exposure so that the therapeutic potential of targeting ERRs *in vivo* could be evaluated. Our work has, for the first time, directly demonstrated the cardioprotective effects of two structurally distinct pan ERR agonists, which strongly supports further efforts in developing additional compounds for preclinical and clinical studies with favorable pharmacodynamic properties and/or isoform specific compounds that potentially optimize the therapeutic effects over the untoward effects.

Our transcriptomic and metabolomics data highlight the activation of the FAO pathway as the main contributor for restoring cardiac function by ERR pan-agonists, as well as activation of autophagy and suppression of cell cycle and developmental genes (Figure 4-6, Supplementary Figure 6). We showed that ERR agonism increased the expression level of *Pdk4* both *in vitro* and *in vivo* (Figure 4D), a well studied ERR-targeted metabolic master regulator, which suppresses glucose ultility and stimulates the FAO pathway in the heart and other metabolically active tissues (57–61). Little is known about the regulatory mechanism of ERR on cell cyle and fetal genes. A previous study on ERRα-null mouse with microarray demonstrated a set of genes involved in cell cycle, proliferation, migration and adhesion were transcriptionally upregulated compared with WT mice (11). Our study unraveled a novel role of E2F1 in mediating the downregulation of cell cycle genes upon ERR agonism (Figure 8). The cardioprotective effects from E2F1 suppression has also been appreciated by previous studies, which showed E2F1 kncokdown reduced mitochondrial fragmentation, cardiomyocyte apoptosis, and myocardial infraction of mouse heart with ischemia/reperfusion injury (66, 67). Mechanistically, ERR may directly regulate *E2f1* transcription level by binding on the promoter region thus altering the transcription of E2F1-dependent cell cycle genes. E2F1 is not likely to be the sole mediator as only partial E2F1 dependancy was observed in the siRNA knockdown study (Figure 8D and Supplementary Figure 9). For instance, A ChIP-on-chip analysis in the same study showed ERRα/γ directly binds to the promoters of cell cycle genes such as *Cenpa* and *Cenpt* (Table 1) (11). Therefore, multiple mediators or even direct suppression through ERR may participate in the transcriptional suppression of cell cycle genes upon ERR agonism. The detailed mechanism requires further investigation.

We found ERR pan-agonists had no effect on cardiac hypertrophy despite being cardioprotective for contractile function (Figure 2). ERR-pan agonists also had no effects in PE-induced cardiomyocytes hypertrophy or activation of ERK1/2 and calcineurin/NFAT signaling pathways in NRVMs (Figure 3). Interestingly, similar results was reported on PPARα, a transcriptional activator of fatty acid metabolic genes in the heart, where cardiac-specific overexpression also led to an elevation in EF without attenuating the cardiac hypertrophy after 8-week TAC (68). Therefore, increasing fatty acid metabolism may provide cardiac contractile function benefit without affecting hypertrophic signaling at least in TAC heart without releaving the pressure overload.

Although ERR agonism appears to change the failing heart to a hypertrophic heart with normal cardiac function, by comparing with the previously reported pathological hypertrophy-specific and physiological hypertrophy-specific gene sets (49), we showed that with treatment of 332, 6-week TAC still induced an expression pattern that more resembles pathological hypertrophy rather than physiological hypertrophy (Figure 3F and 3G). This data indicate that the signal for initiating the pathological hypertrophy is triggered very early after the onset of TAC, and the process is irreversible by metabolic remodeling even when the cardiac function is preserved. However, with sufficient energy supply, the transition from adaptive cardiac hypertrophy to malapative pump failure may not occur and a normal cardiac function can be maintained even in the presence of hypertrophy and persistently increased hemodynamic load. This is critical for most patients with heart failure, who already have significant cardiac remodeling due to long standing disease.

PPARα is also an activator of fatty acid metabolism, which has been proposed as therapeutic target for heart failure (69). However, clinical use of PPAR agonists has shown to increase adverse cardiovascular events (nonfatal MI, transient ischemic attack, congestive heart failure, etc.) or even worsen heart failure (22, 70–72). Several mechanisms may contribute to the adverse effects. First, PPARα activates genes for fatty acid uptake and β-oxidation but suppresses mitochondrial OXPHOS genes (73, 74), which leads to lipid accumulation and toxicity. A previous study demonstrated that after a short-term fast which activates PPARα in the heart and liver (75, 76), a remarkable accumulation of lipid droplets stained with oil red O were observed in cardiac myocytes sections from a PPARα-overexpression mouse compared with WT (73). In contrast, ERR agonists improve overall metabolic profiling in the heart and is of limited concern of lipotoxicity (77). Indeed, we did not see accumulation of lipid droplets in the EM performed after 6 weeks of ERR agonist administration (Figure 1D). Second, mouse studies suggest that the effects of PPARα activation depend on the phase of heart failure. At the early phase of heart failure, the hypertrophic heart undergoes substrate switch and becomes more reliant on glucose oxidation but not FAO; treatment with PPARα agonists at this stage is actually harmful (78), whereas PPARα activation during the progression of heart failure improves cardiac function (68). Compared to PPAR, ERR not only upregulates fatty acid metabolism but also a broader spectrum of other metabolic pathways including glucose metabolism and OXPHOS (Figure 4-5, Supplementary Figure 5-7). Therefore, we speculate that ERR activation could be beneficial to patients at any stage of heart failure as the metabolic plasticity is preserved and enhanced. Third, some PPAR agonist side effects observed in animal studies might be due to off-target effects. For instance, the PPARα agonist fenofibrate was found to exacerbate LV dilation and cardiac fibrosis in PPARα-knockout mice with pressure overload (79). We did not notice any overt side effects from 332 or 915 treatment *in vivo*, although more detailed study is required.

Acute knockdown of individual ERR isoforms *in vitro* confirmed the ERR dependency of 332/915-induced transcriptional regulation (Figure 7 and Supplementary Figure 8). Given that ERRα and ERRγ share many common target genes (11), it is somewhat surprising that ERRγ, but not ERRα, is responsible for most of the 332/915-induced metabolic gene transcription. Previous studies showed that all ERR isoforms exhibit a basal level of transcriptional activity which is independent of ligand binding (80–85). Different ERR isoforms may have different levels of basal activity resulting in different inducible activity by ERR agonist. Moreover, ERR isoforms can function as monomers, homodimers, or heterodimers, which modulates the gene recognition pattern and transcriptional activity (84, 86–89). For instance, homodimerization enhances the transcriptional activity of ERRγ, while heterodimerization of ERRα and ERRγ suppresses the transcriptional activity of ERRα or ERRγ alone (87). Induction of *Esrrb* and *Esrrg* transcription was also observed after 332 and 915 administration (Figure 4C), which suggests a positive feedback regulatory mechanism upon activation of ERR and further complicates the isoform-specific regulation. Thus, the stoichiometric ratio of each individual ERR isoform is likely to affect the pharmacological outcome upon ERR pan-agonist stimulation. Further study with isoform-specific agonists combined with isoform-specific mutant mouse models is required to address all these possibilities.

In summary, our study demonstrated that activation of ERRs by ERR pan-agonists 332 or 915 protects cardiac function against heart failure predominently through elevating mitochondrial function and boosting fatty acid metabolism. ERR agonism also leads to suppression of cell cycle genes transcription which is partially mediated by E2F1. ERRγ may be the main mediator for transcriptional regulation induced by 332 or 915 in cardiomyocytes. Thus, ERR pan-agonists may be a viable therapeutic strategy for improving metabolic gene profiling and pump function in heart failure.

## Author Contributions

LZ and TB conceived the project. LZ, TB, CB, and WX designed the experiments. WX, CB, HL, MH, KY, ML, CH, CA, AG, CF, and RW performed the experiments and analyzed the data. WX, CB, KY, EN, RE, LP, JW, AM, TB, and LZ wrote the manuscript.

## Acknowledgments

The work is supported by NIH grant K08 HL123551 (LZ), and R01 HL143067 (LZ).

---

**Supplementary Figure 1**

**Echocardiography measurements for assessing the effects of ERR agonists in TAC-induced pressure-overload model.**

**(A)** Fraction shortening (FS). **(B)** Left ventricular end-systolic posterior wall thickness (LVPW;s). **(C)** Left ventricular end-systolic anterior wall thickness (LVAW;s). **(D)** Left ventricular mass calculated based on anterior wall (LV Mass AW). **(E)** Left ventricular end-diastolic diameter (LVID;d). **(F)** Left ventricular end-systolic diameter (LVID;s) **(G)** Left ventricular end-diastolic volume (LV Vol;d), **(H)** Left ventricular end-systolic volume (LV Vol;s), **(I)** Stroke volume (SV). **(J)** Body weight. **(K)** Ratio of lung weight to body weight. N=4-10. Statisctic analysis was performed for data in (K) using two-way ANOVA. Multiple comparison is corrected by Tukey method with α=0.05. Data are presented as mean ± SEM.

**Supplementary Figure 2**

**Quantification of mitochondrial gene copy number upon 332 treatment.**

(A) Mitochondrial gene copy numbers measured from mouse heart samples. N=4-6. *Chr6* was used as internal control. (B) Mitochondrial gene copy numbers measured from NRVMs. N=3. *Chr4* was used as internal control. Statistical analysis was performed using one-way ANOVA. Multiple comparison is corrected by Dunnet method with α=0.05. Data are presented as mean ± SEM.

**Supplementary Figure 3**

**Examination of the specificity of SLU-PP-915.**

**(A)** Results of full length ERRα, ERRb or ERRg co-transfection assays in HEK293 cells illustrating the specific activity of SLU-PP-915 (n=4). **(B)** Specificity of 915 (10μM) on a complete panel of Gal4-DDB NR-LBD chimeric nuclear receptor in luciferase reporter assay. Data are presented as relative to vehicle treated cells and some activity was noted at other nuclear receptors. Student’s test was used to calculate statistical significance. P<0.05 was considered significant. **(C)** Full dose response of 915 or ligand control in the same luciferase assay for positive hits identified in (B). **(D)** 332 and 915 specificity in the NIH PDSP (Psychoactive Drug Screening Program) against a panel of GPCRs, ion channels and transporters in a radioligand binding assay format. Data are presented as mean ± SEM.

**Supplementary Figure 4**

**Pharmacokinetic analysis of 915.**

Males (n=3 per time point) C57Bl/6J mice (8 weeks of age at study initiation) were dosing intra-peritoneally with SLU-PP-915 (20mg/kg, 10% DMSO, 10% tween in PBS). Blood was collected at different time point and plasma was analyzed for 915 quantification using LC-MS/MS. Table shows the summary of pharmacokinetic properties. Data represented as mean and are generated by ADME Solutions, Inc.

**Supplementary Figure 5**

**Metabolic and cell cycle-related pathways were significantly altered upon ERR agonism.**

**(A)** and **(B)** Pathway enrichment analysis of RNAseq of NRVM treated with vehicle, 10μM 332, or 10μM 915 for 72hr. Genes with 1.5-fold increase versus vehicle group and adjusted p-value<0.05 (DEGs) were included in the analysis. Gene ontology biological processes (GO BP) term enrichment analysis and KEGG pathway analysis are shown in (A) and (B). Pathway enrichment with FDR (false discovery rate) lower than 0.05 was considered as statistically significant.

**Supplementary Figure 6**

**Deconvolution of mouse heart bulk RNA-seq revealed ERR agonism upregulated OXPHOS and downregulated Hippo signaling pathway in cardiomyocyte *in vivo*.**

**(A)** XDec deconvolution identified 9 cell types in vehicle or 332 treated TAC mice. 3 biological replicates of mouse heart tissues were used for deconvolution. Reference profiles of cardiomyocyte, endothelial cell, fibroblast, and immune cells from previously published single-cell RNA sequencing data (GSE120064) were indicated in red, brown, green, and blue. Distribution of correlation between the expression profiles of reference cell type and deconvoluted constituent cell type is shown on the right top panel. See details in Method. **(B)** Proportions of CM profiles in Sham+Veh, Sham+332, TAC+Veh, and TAC+332 whole heart samples (n=3). **(C)** Proportions of CM profile 2, 3 and 4 in TAC+Veh and TAC+332 samples. Welch Two Sample t-test was used to determine statistical difference. **(D)** KEGG pathway analysis of upregulated and downregulated DEGs between CM profile 3 and 4.Pathway enrichment with p-value<0.05 was considered as statistically significant.

**Supplementary Figure 7**

**ERR agonism upregulated genes involved in TCA cycle and glycolysis *in vitro*.**

**(A)** Heatmap of genes involved in TCA cycle and upregulated by 332 and 915. N=3. **(B)** Heatmap of genes involved in glycolysis and upregulated by 332 and 915. N=3. Genes with significant changes were included (adjust p-value<0.05).

**Supplementary Figure 8**

**Effects of ERR isoform knockdown on ERR agonist-induced gene expressions.**

(A) and (B) Metabolic genes and Autophgy genes. MNRVMs were transfected with 25pM scramble or ERR siRNA. 24hr after transfection, NRVMs were treated with vehicle, 10μM 332, or 10μM 915 for 24hr. Gene expression level was quantified by qPCR and normalized to the vehicle scramble group. N=3-6. *p<0.05 vs vehicle within each transfection group. ^#^p<0.05 vs scramble with the same treatment (data points with the same color). Statistical differences were determined by two-way ANOVA. Multiple comparison is corrected by Tukey method with α=0.05. Data are presented as mean ± SEM.

**Supplementary Figure 9**

**E2F1-independent DEGs.**

Downregulated DEGs with predicted E2F1 cognate site and over 1.5-fold change in expression upon ERR agonism were selected and examined in the experiment. Statistical differences were determined by two-way ANOVA. Multiple comparison is corrected by Tukey method with α=0.05. Data are presented as mean ± SEM.

## Reference

1. Murphy SP, Ibrahim NE, and Januzzi JL, Jr. Heart Failure With Reduced Ejection Fraction: A Review. JAMA. 2020;324(5):488–504.

2. Heidenreich PA, Albert NM, Allen LA, Bluemke DA, Butler J, Fonarow GC, et al. Forecasting the impact of heart failure in the United States: a policy statement from the American Heart Association. Circ Heart Fail. 2013;6(3):606–19.

3. Mazurek JA, and Jessup M. Understanding Heart Failure. Heart Fail Clin. 2017;13(1):1–19.

4. Taylor CJ, Ordonez-Mena JM, Roalfe AK, Lay-Flurrie S, Jones NR, Marshall T, et al. Trends in survival after a diagnosis of heart failure in the United Kingdom 2000-2017: population based cohort study. BMJ. 2019;364:l223.

5. Stanley WC, Recchia FA, and Lopaschuk GD. Myocardial substrate metabolism in the normal and failing heart. Physiol Rev. 2005;85(3):1093–129.

6. Zhou B, and Tian R. Mitochondrial dysfunction in pathophysiology of heart failure. J Clin Invest. 2018;128(9):3716–26.

7. Bertero E, and Maack C. Metabolic remodelling in heart failure. Nat Rev Cardiol. 2018;15(8):457–70.

8. Noordali H, Loudon BL, Frenneaux MP, and Madhani M. Cardiac metabolism - A promising therapeutic target for heart failure. Pharmacol Ther. 2018;182:95–114.

9. Huss JM, Imahashi K, Dufour CR, Weinheimer CJ, Courtois M, Kovacs A, et al. The nuclear receptor ERRalpha is required for the bioenergetic and functional adaptation to cardiac pressure overload. Cell Metab. 2007;6(1):25–37.

10. Wang T, McDonald C, Petrenko NB, Leblanc M, Wang T, Giguere V, et al. Estrogen-related receptor alpha (ERRalpha) and ERRgamma are essential coordinators of cardiac metabolism and function. Mol Cell Biol. 2015;35(7):1281–98.

11. Dufour CR, Wilson BJ, Huss JM, Kelly DP, Alaynick WA, Downes M, et al. Genome-wide orchestration of cardiac functions by the orphan nuclear receptors ERRalpha and gamma. Cell Metab. 2007;5(5):345–56.

12. Alaynick WA, Kondo RP, Xie W, He W, Dufour CR, Downes M, et al. ERRgamma directs and maintains the transition to oxidative metabolism in the postnatal heart. Cell Metab. 2007;6(1):13–24.

13. Sakamoto T, Matsuura TR, Wan S, Ryba DM, Kim JU, Won KJ, et al. A Critical Role for Estrogen-Related Receptor Signaling in Cardiac Maturation. Circ Res. 2020;126(12):1685–702.

14. Sihag S, Cresci S, Li AY, Sucharov CC, and Lehman JJ. PGC-1alpha and ERRalpha target gene downregulation is a signature of the failing human heart. J Mol Cell Cardiol. 2009;46(2):201–12.

15. Schilling J, and Kelly DP. The PGC-1 cascade as a therapeutic target for heart failure. J Mol Cell Cardiol. 2011;51(4):578–83.

16. Vega RB, and Kelly DP. Cardiac nuclear receptors: architects of mitochondrial structure and function. J Clin Invest. 2017;127(4):1155–64.

17. Suetsugi M, Su L, Karlsberg K, Yuan YC, and Chen S. Flavone and isoflavone phytoestrogens are agonists of estrogen-related receptors. Mol Cancer Res. 2003;1(13):981–91.

18. Lynch C, Zhao J, Huang R, Kanaya N, Bernal L, Hsieh JH, et al. Identification of Estrogen-Related Receptor alpha Agonists in the Tox21 Compound Library. Endocrinology. 2018;159(2):744–53.

19. Fujiwara Y, Deguchi K, Naka Y, Sasaki M, Nishimoto T, and Yoshida Y. Development of matured hiPSCs-derived 3D cardiac tissue using ERR gamma agonist and mechanical stress and application for Hypertrophic Cardiomyopathy (HCM) model. European Heart Journal. 2020;41(Supplement_2).

20. Zuercher WJ, Gaillard S, Orband-Miller LA, Chao EY, Shearer BG, Jones DG, et al. Identification and structure-activity relationship of phenolic acyl hydrazones as selective agonists for the estrogen-related orphan nuclear receptors ERRbeta and ERRgamma. J Med Chem. 2005;48(9):3107–9.

21. Wang L, Zuercher WJ, Consler TG, Lambert MH, Miller AB, Orband-Miller LA, et al. X-ray crystal structures of the estrogen-related receptor-gamma ligand binding domain in three functional states reveal the molecular basis of small molecule regulation. J Biol Chem. 2006;281(49):37773–81.

22. Nissen SE, Wolski K, and Topol EJ. Effect of muraglitazar on death and major adverse cardiovascular events in patients with type 2 diabetes mellitus. JAMA. 2005;294(20):2581–6.

23. Zhang L, Zhang R, Tien CL, Chan RE, Sugi K, Fu C, et al. REV-ERBalpha ameliorates heart failure through transcription repression. JCI Insight. 2017;2(17).

24. O’Connell TD, Rodrigo MC, and Simpson PC. Isolation and culture of adult mouse cardiac myocytes. Methods Mol Biol. 2007;357:271–96.

25. Collaboration A, Aad G, Abbott B, Abdallah J, Khalek SA, Abdinov O, et al. Search for direct top squark pair production in events with a [Formula: see text] boson, [Formula: see text]-jets and missing transverse momentum in [Formula: see text] TeV [Formula: see text] collisions with the ATLAS detector. Eur Phys J C Part Fields. 2014;74(6):2883.

26. Wu D, Jian C, Peng Q, Hou T, Wu K, Shang B, et al. Prohibitin 2 deficiency impairs cardiac fatty acid oxidation and causes heart failure. Cell Death Dis. 2020;11(3):181.

27. Readnower RD, Brainard RE, Hill BG, and Jones SP. Standardized bioenergetic profiling of adult mouse cardiomyocytes. Physiol Genomics. 2012;44(24):1208–13.

28. Wettmarshausen J, and Perocchi F. In: Raffaello A, and Vecellio Reane D eds. Calcium Signalling: Methods and Protocols. New York, NY: Springer New York; 2019:197–222.

29. Liao X, Zhang R, Lu Y, Prosdocimo DA, Sangwung P, Zhang L, et al. Kruppel-like factor 4 is critical for transcriptional control of cardiac mitochondrial homeostasis. J Clin Invest. 2015;125(9):3461–76.

30. Ge SX, Jung D, and Yao R. ShinyGO: a graphical gene-set enrichment tool for animals and plants. Bioinformatics. 2020;36(8):2628–9.

31. Hulsen T, de Vlieg J, and Alkema W. BioVenn - a web application for the comparison and visualization of biological lists using area-proportional Venn diagrams. BMC Genomics. 2008;9:488.

32. Ren Z, Yu P, Li D, Li Z, Liao Y, Wang Y, et al. Single-Cell Reconstruction of Progression Trajectory Reveals Intervention Principles in Pathological Cardiac Hypertrophy. Circulation. 2020;141(21):1704–19.

33. Lun AT, Bach K, and Marioni JC. Pooling across cells to normalize single-cell RNA sequencing data with many zero counts. Genome Biol. 2016;17:75.

34. Huber W, Carey VJ, Gentleman R, Anders S, Carlson M, Carvalho BS, et al. Orchestrating high-throughput genomic analysis with Bioconductor. Nat Methods. 2015;12(2):115–21.

35. Kanehisa M, and Goto S. KEGG: kyoto encyclopedia of genes and genomes. Nucleic Acids Res. 2000;28(1):27–30.

36. Gelinas R, Mailleux F, Dontaine J, Bultot L, Demeulder B, Ginion A, et al. AMPK activation counteracts cardiac hypertrophy by reducing O-GlcNAcylation. Nat Commun. 2018;9(1):374.

37. Chan AY, Dolinsky VW, Soltys CL, Viollet B, Baksh S, Light PE, et al. Resveratrol inhibits cardiac hypertrophy via AMP-activated protein kinase and Akt. J Biol Chem. 2008;283(35):24194–201.

38. deAlmeida AC, van Oort RJ, and Wehrens XH. Transverse aortic constriction in mice. J Vis Exp. 2010(38).

39. Chaanine AH, Joyce LD, Stulak JM, Maltais S, Joyce DL, Dearani JA, et al. Mitochondrial Morphology, Dynamics, and Function in Human Pressure Overload or Ischemic Heart Disease With Preserved or Reduced Ejection Fraction. Circ Heart Fail. 2019;12(2):e005131.

40. Heineke J, and Molkentin JD. Regulation of cardiac hypertrophy by intracellular signalling pathways. Nat Rev Mol Cell Biol. 2006;7(8):589–600.

41. Schiattarella GG, Hill TM, and Hill JA. Is Load-Induced Ventricular Hypertrophy Ever Compensatory? Circulation. 2017;136(14):1273–5.

42. Molkentin JD, Lu JR, Antos CL, Markham B, Richardson J, Robbins J, et al. A calcineurin-dependent transcriptional pathway for cardiac hypertrophy. Cell. 1998;93(2):215–28.

43. Bueno OF, De Windt LJ, Tymitz KM, Witt SA, Kimball TR, Klevitsky R, et al. The MEK1-ERK1/2 signaling pathway promotes compensated cardiac hypertrophy in transgenic mice. EMBO J. 2000;19(23):6341–50.

44. Nakamura M, and Sadoshima J. Mechanisms of physiological and pathological cardiac hypertrophy. Nat Rev Cardiol. 2018;15(7):387–407.

45. Davis J, Davis LC, Correll RN, Makarewich CA, Schwanekamp JA, Moussavi-Harami F, et al. A Tension-Based Model Distinguishes Hypertrophic versus Dilated Cardiomyopathy. Cell. 2016;165(5):1147–59.

46. Gineitis D, and Treisman R. Differential usage of signal transduction pathways defines two types of serum response factor target gene. J Biol Chem. 2001;276(27):24531–9.

47. Goetze S, Kintscher U, Kaneshiro K, Meehan WP, Collins A, Fleck E, et al. TNFalpha induces expression of transcription factors c-fos, Egr-1, and Ets-1 in vascular lesions through extracellular signal-regulated kinases 1/2. Atherosclerosis. 2001;159(1):93–101.

48. Khachigian LM. Early growth response-1 in cardiovascular pathobiology. Circ Res. 2006;98(2):186–91.

49. Bostrom P, Mann N, Wu J, Quintero PA, Plovie ER, Panakova D, et al. C/EBPbeta controls exercise-induced cardiac growth and protects against pathological cardiac remodeling. Cell. 2010;143(7):1072–83.

50. Diwan A, and Dorn GW, 2nd. Decompensation of cardiac hypertrophy: cellular mechanisms and novel therapeutic targets. Physiology (Bethesda). 2007;22:56–64.

51. Mia MM, and Singh MK. The Hippo Signaling Pathway in Cardiac Development and Diseases. Front Cell Dev Biol. 2019;7:211.

52. Zhang Y, Mignone J, and MacLellan WR. Cardiac Regeneration and Stem Cells. Physiol Rev. 2015;95(4):1189–204.

53. Lee HG, Chen Q, Wolfram JA, Richardson SL, Liner A, Siedlak SL, et al. Cell cycle re-entry and mitochondrial defects in myc-mediated hypertrophic cardiomyopathy and heart failure. PLoS One. 2009;4(9):e7172.

54. Pereira AHM, Cardoso AC, Consonni SR, Oliveira RR, Saito A, Vaggione MLB, et al. MEF2C repressor variant deregulation leads to cell cycle re-entry and development of heart failure. EBioMedicine. 2020;51:102571.

55. Cui M, Wang Z, Bassel-Duby R, and Olson EN. Genetic and epigenetic regulation of cardiomyocytes in development, regeneration and disease. Development. 2018;145(24).

56. Onuchic V, Hartmaier RJ, Boone DN, Samuels ML, Patel RY, White WM, et al. Epigenomic Deconvolution of Breast Tumors Reveals Metabolic Coupling between Constituent Cell Types. Cell Rep. 2016;17(8):2075–86.

57. Wende AR, Huss JM, Schaeffer PJ, Giguere V, and Kelly DP. PGC-1alpha coactivates PDK4 gene expression via the orphan nuclear receptor ERRalpha: a mechanism for transcriptional control of muscle glucose metabolism. Mol Cell Biol. 2005;25(24):10684–94.

58. Zhang Y, Ma K, Sadana P, Chowdhury F, Gaillard S, Wang F, et al. Estrogen-related receptors stimulate pyruvate dehydrogenase kinase isoform 4 gene expression. J Biol Chem. 2006;281(52):39897–906.

59. Araki M, and Motojima K. Identification of ERRalpha as a specific partner of PGC-1alpha for the activation of PDK4 gene expression in muscle. FEBS J. 2006;273(8):1669–80.

60. Zhang S, Hulver MW, McMillan RP, Cline MA, and Gilbert ER. The pivotal role of pyruvate dehydrogenase kinases in metabolic flexibility. Nutr Metab (Lond). 2014;11(1):10.

61. Zhao G, Jeoung NH, Burgess SC, Rosaaen-Stowe KA, Inagaki T, Latif S, et al. Overexpression of pyruvate dehydrogenase kinase 4 in heart perturbs metabolism and exacerbates calcineurin-induced cardiomyopathy. Am J Physiol Heart Circ Physiol. 2008;294(2):H936–43.

62. Orogo AM, and Gustafsson AB. Therapeutic targeting of autophagy: potential and concerns in treating cardiovascular disease. Circ Res. 2015;116(3):489–503.

63. Yamamoto A, Tagawa Y, Yoshimori T, Moriyama Y, Masaki R, and Tashiro Y. Bafilomycin A1 prevents maturation of autophagic vacuoles by inhibiting fusion between autophagosomes and lysosomes in rat hepatoma cell line, H-4-II-E cells. Cell Struct Funct. 1998;23(1):33–42.

64. Rowe GC, Asimaki A, Graham EL, Martin KD, Margulies KB, Das S, et al. Development of dilated cardiomyopathy and impaired calcium homeostasis with cardiac-specific deletion of ESRRbeta. Am J Physiol Heart Circ Physiol. 2017;312(4):H662–H71.

65. Kent LN, and Leone G. The broken cycle: E2F dysfunction in cancer. Nat Rev Cancer. 2019;19(6):326–38.

66. Wang K, Zhou LY, Wang JX, Wang Y, Sun T, Zhao B, et al. E2F1-dependent miR-421 regulates mitochondrial fragmentation and myocardial infarction by targeting Pink1. Nat Commun. 2015;6:7619.

67. Wang K, An T, Zhou LY, Liu CY, Zhang XJ, Feng C, et al. E2F1-regulated miR-30b suppresses Cyclophilin D and protects heart from ischemia/reperfusion injury and necrotic cell death. Cell Death Differ. 2015;22(5):743–54.

68. Kaimoto S, Hoshino A, Ariyoshi M, Okawa Y, Tateishi S, Ono K, et al. Activation of PPAR-alpha in the early stage of heart failure maintained myocardial function and energetics in pressure-overload heart failure. Am J Physiol Heart Circ Physiol. 2017;312(2):H305–H13.

69. Sarma S, Ardehali H, and Gheorghiade M. Enhancing the metabolic substrate: PPAR-alpha agonists in heart failure. Heart Fail Rev. 2012;17(1):35–43.

70. Lipscombe J, Lewis GF, Cattran D, and Bargman JM. Deterioration in renal function associated with fibrate therapy. Clin Nephrol. 2001;55(1):39–44.

71. Erdmann E, and Wilcox RG. Weighing up the cardiovascular benefits of thiazolidinedione therapy: the impact of increased risk of heart failure. Eur Heart J. 2008;29(1):12–20.

72. Sarma S. Use of clinically available PPAR agonists for heart failure; do the risks outweigh the potential benefits? Curr Mol Pharmacol. 2012;5(2):255–63.

73. Finck BN, Lehman JJ, Leone TC, Welch MJ, Bennett MJ, Kovacs A, et al. The cardiac phenotype induced by PPARalpha overexpression mimics that caused by diabetes mellitus. J Clin Invest. 2002;109(1):121–30.

74. Oka S, Alcendor R, Zhai P, Park JY, Shao D, Cho J, et al. PPARalpha-Sirt1 complex mediates cardiac hypertrophy and failure through suppression of the ERR transcriptional pathway. Cell Metab. 2011;14(5):598–611.

75. Leone TC, Weinheimer CJ, and Kelly DP. A critical role for the peroxisome proliferator-activated receptor alpha (PPARalpha) in the cellular fasting response: the PPARalpha-null mouse as a model of fatty acid oxidation disorders. Proc Natl Acad Sci U S A. 1999;96(13):7473–8.

76. Kersten S, Seydoux J, Peters JM, Gonzalez FJ, Desvergne B, and Wahli W. Peroxisome proliferator-activated receptor alpha mediates the adaptive response to fasting. J Clin Invest. 1999;103(11):1489–98.

77. Fan W, and Evans R. PPARs and ERRs: molecular mediators of mitochondrial metabolism. Curr Opin Cell Biol. 2015;33:49–54.

78. Young ME, Laws FA, Goodwin GW, and Taegtmeyer H. Reactivation of peroxisome proliferator-activated receptor alpha is associated with contractile dysfunction in hypertrophied rat heart. J Biol Chem. 2001;276(48):44390–5.

79. Duhaney TA, Cui L, Rude MK, Lebrasseur NK, Ngoy S, De Silva DS, et al. Peroxisome proliferator-activated receptor alpha-independent actions of fenofibrate exacerbates left ventricular dilation and fibrosis in chronic pressure overload. Hypertension. 2007;49(5):1084–94.

80. Xie W, Hong H, Yang NN, Lin RJ, Simon CM, Stallcup MR, et al. Constitutive activation of transcription and binding of coactivator by estrogen-related receptors 1 and 2. Mol Endocrinol. 1999;13(12):2151–62.

81. Hong H, Yang L, and Stallcup MR. Hormone-independent transcriptional activation and coactivator binding by novel orphan nuclear receptor ERR3. J Biol Chem. 1999;274(32):22618–26.

82. Chen S, Zhou D, Yang C, and Sherman M. Molecular basis for the constitutive activity of estrogen-related receptor alpha-1. J Biol Chem. 2001;276(30):28465–70.

83. Kallen J, Schlaeppi JM, Bitsch F, Filipuzzi I, Schilb A, Riou V, et al. Evidence for ligand-independent transcriptional activation of the human estrogen-related receptor alpha (ERRalpha): crystal structure of ERRalpha ligand binding domain in complex with peroxisome proliferator-activated receptor coactivator-1alpha. J Biol Chem. 2004;279(47):49330–7.

84. Huss JM, Garbacz WG, and Xie W. Constitutive activities of estrogen-related receptors: Transcriptional regulation of metabolism by the ERR pathways in health and disease. Biochim Biophys Acta. 2015;1852(9):1912–27.

85. Greschik H, Wurtz JM, Sanglier S, Bourguet W, van Dorsselaer A, Moras D, et al. Structural and functional evidence for ligand-independent transcriptional activation by the estrogen-related receptor 3. Mol Cell. 2002;9(2):303–13.

86. Gearhart MD, Holmbeck SM, Evans RM, Dyson HJ, and Wright PE. Monomeric complex of human orphan estrogen related receptor-2 with DNA: a pseudo-dimer interface mediates extended half-site recognition. J Mol Biol. 2003;327(4):819–32.

87. Huppunen J, and Aarnisalo P. Dimerization modulates the activity of the orphan nuclear receptor ERRgamma. Biochem Biophys Res Commun. 2004;314(4):964–70.

88. Huppunen J, Wohlfahrt G, and Aarnisalo P. Requirements for transcriptional regulation by the orphan nuclear receptor ERRgamma. Mol Cell Endocrinol. 2004;219(1-2):151–60.

89. Zhang Z, and Teng CT. Interplay between estrogen-related receptor alpha (ERRalpha) and gamma (ERRgamma) on the regulation of ERRalpha gene expression. Mol Cell Endocrinol. 2007;264(1-2):128–41.

