## supplemental for "Novel ERR pan-agonists ameliorate heart failure through boosting cardiac fatty acid metabolism and mitochondrial function"

### Supplementary Figure 1

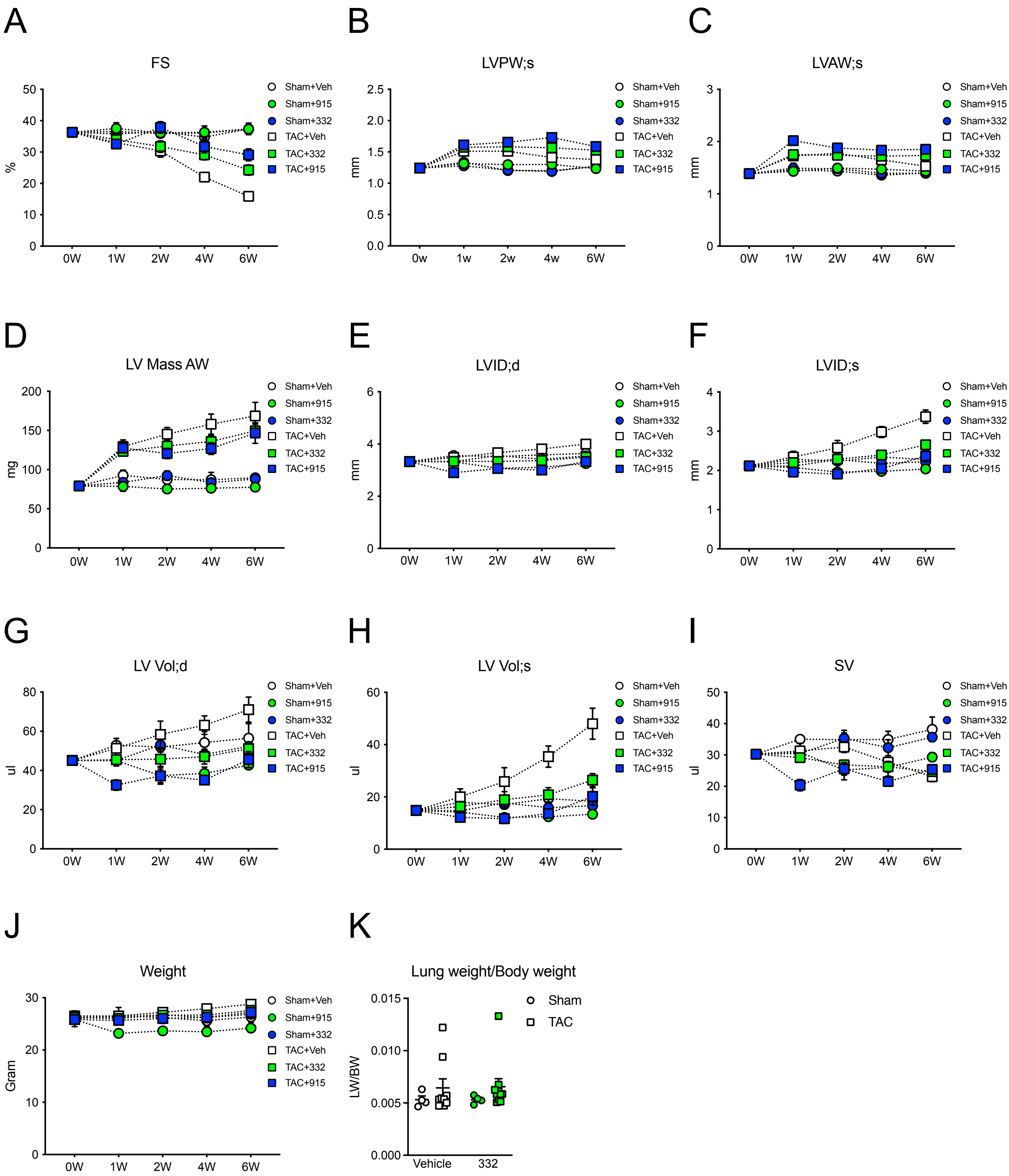

Supplementary Figure 2

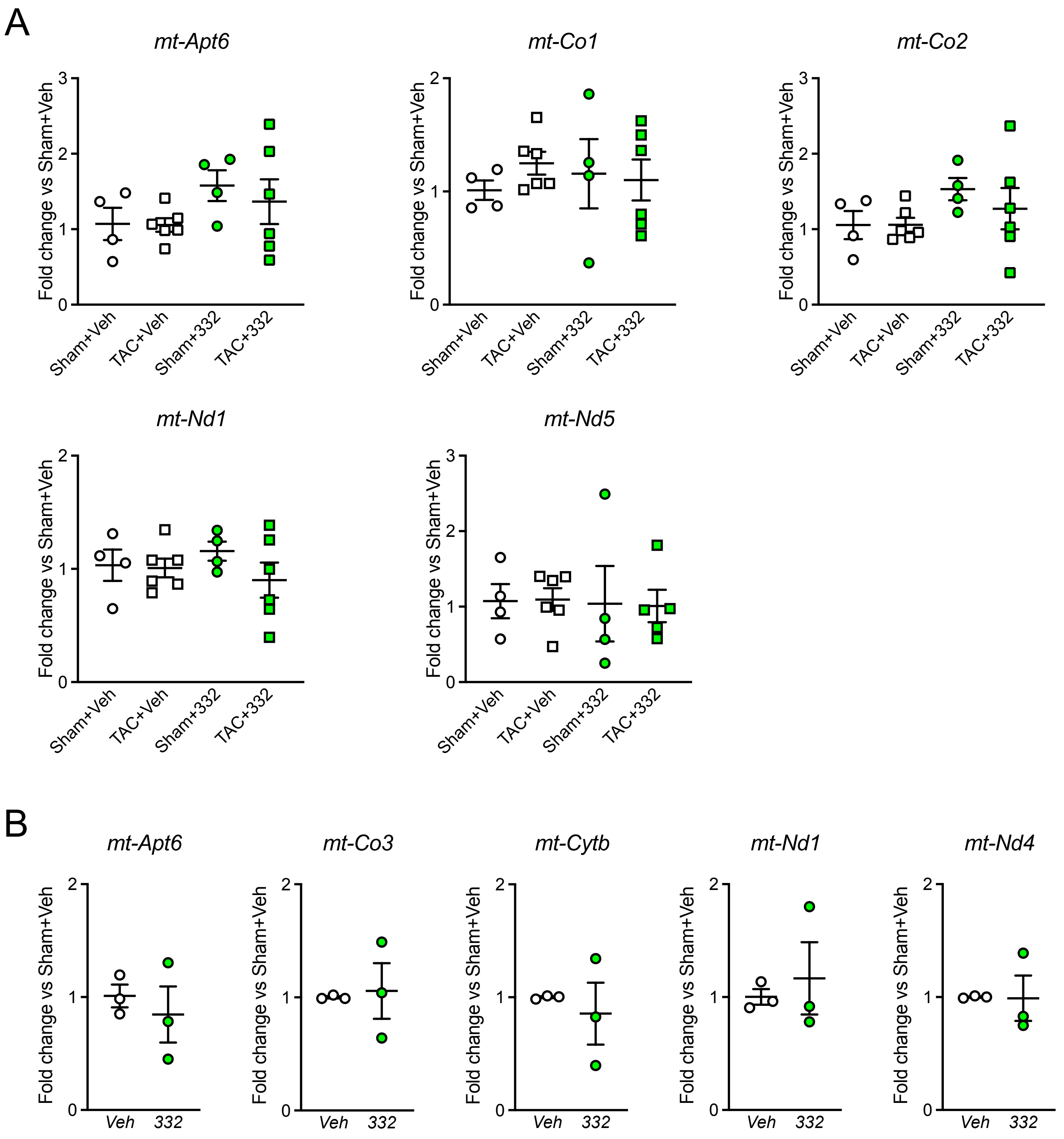

### Supplementary Figure 3

A

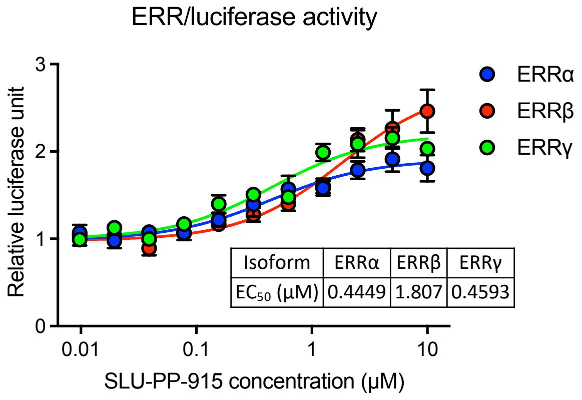


B

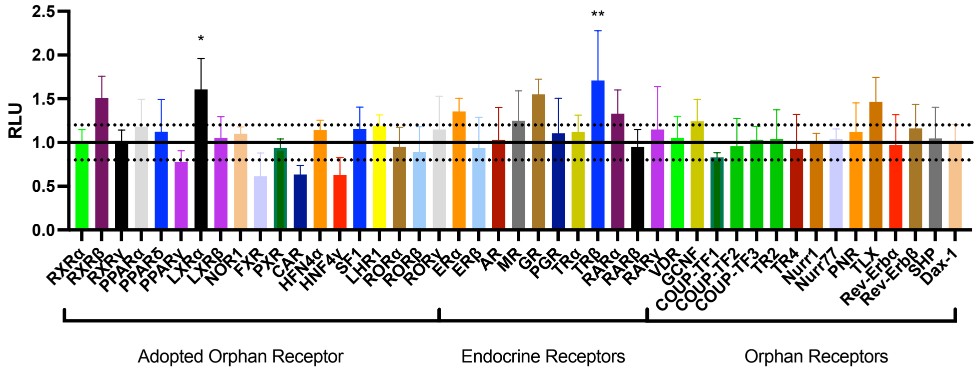


C

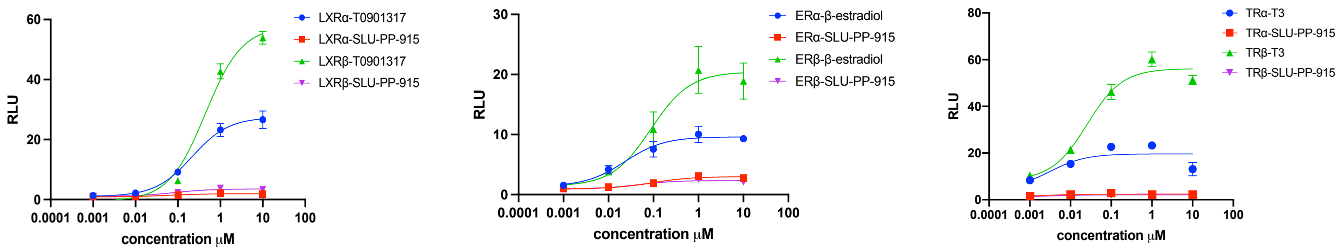


D

| Ligands | Receptors tested | EC50 |
|---|---|---|
| SLU-PP-332 | 5-HT1A, 5-HT1B, 5-HT1D, 5-HT1E, 5-HT2A, 5-HT3, 5-HT5A, 5-HT6, 5-HT7A, Alpha1A, Alpha1B, Alpha1D, Alpha2A, Alpha2B, Alpha2C, Beta1, Beta2, Beta3, BZP Rat Brain Site, D1, D2, D3, D4, D5, DAT, DOR, GABAA, H1, H2, H3, H4, KOR, M1, M2, M3, M4, M5, MOR, NET, PBR, SERT, Sigma 1, Sigma 2 | >10μM |
| SLU-PP-915 | 5-HT1A, 5-HT1B, 5-HT1D, 5-HT1E, 5-HT2A, 5-HT2B, 5-HT2C, 5-HT3, 5-HT5A, 5-HT7A, Alpha1A, Alpha1B, Alpha1D, Alpha2A, Alpha2B, Alpha2C, Beta1, Beta2, Beta3, BZP Rat Brain Site, D1, D2, D3, D4, D5, DAT, DOR, GABAA, H1, H2, H3, H4, KOR, M1, M2, M3, M4, M5, MOR, NET, PBR, SERT, Sigma 1, Sigma 2 | >10μM |

Supplementary Figure 4

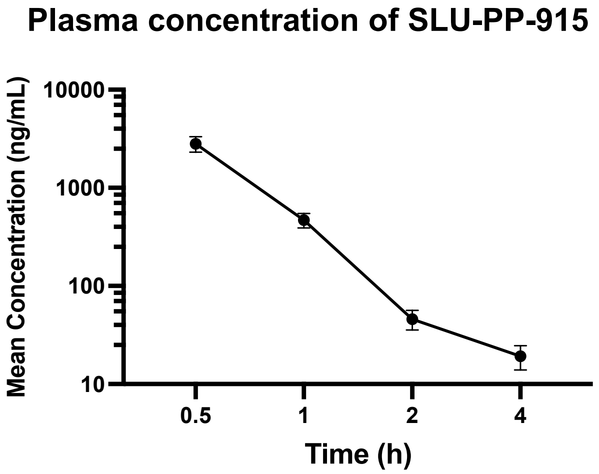


| Compound | Tmax (h) | Cmax (ng/mL) | AUClast (h*ng/mL) | Vz/F (L/kg) | CL/F (mL/min/kg) | $t_{1/2}$ (h) |
|---|---|---|---|---|---|---|
| SLU-PP-915 | 0.5 | 2813 | 1846 | 11 | 179 | 0.71 |

Supplementary Figure 5

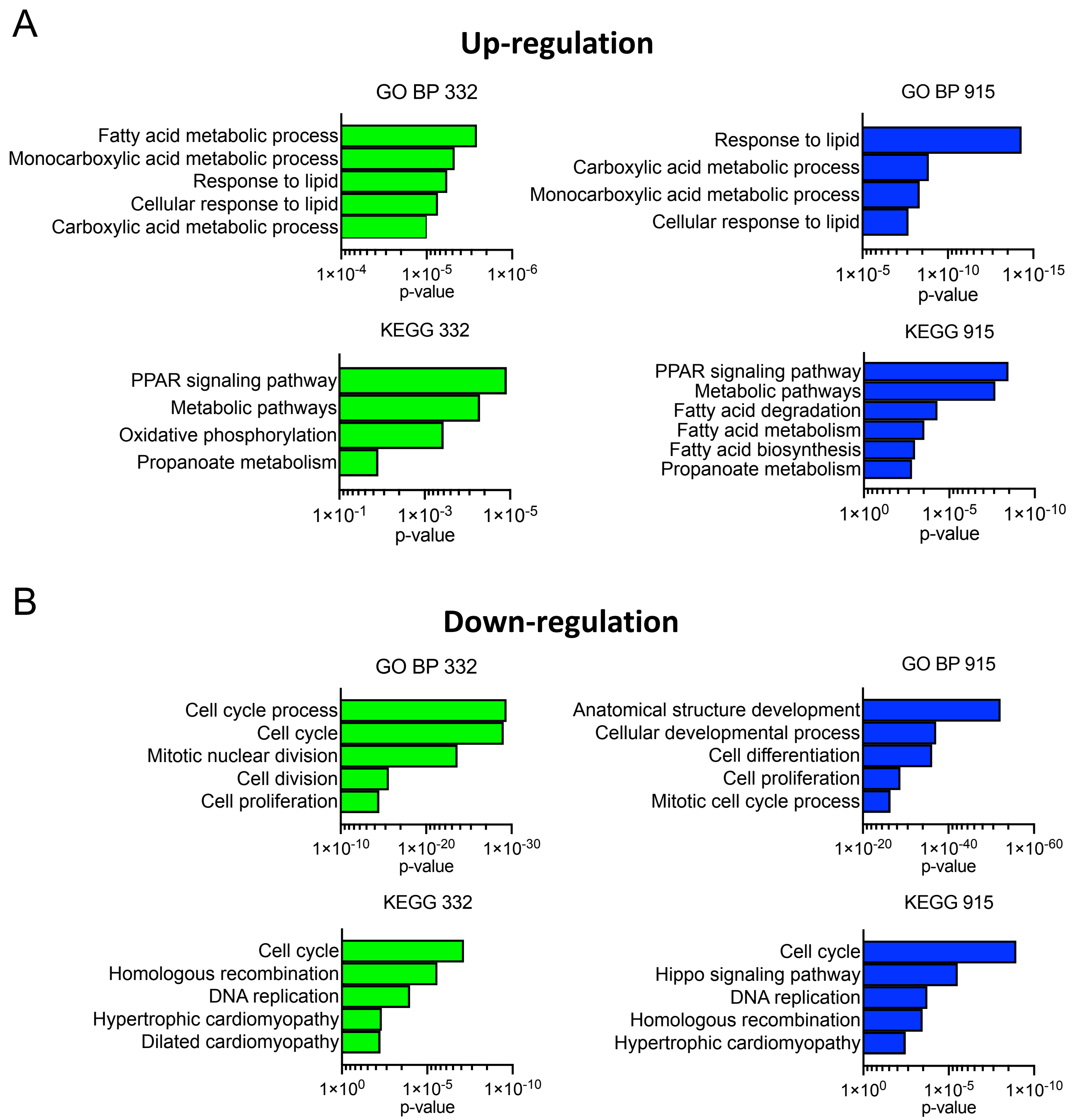

### Supplementary Figure 6

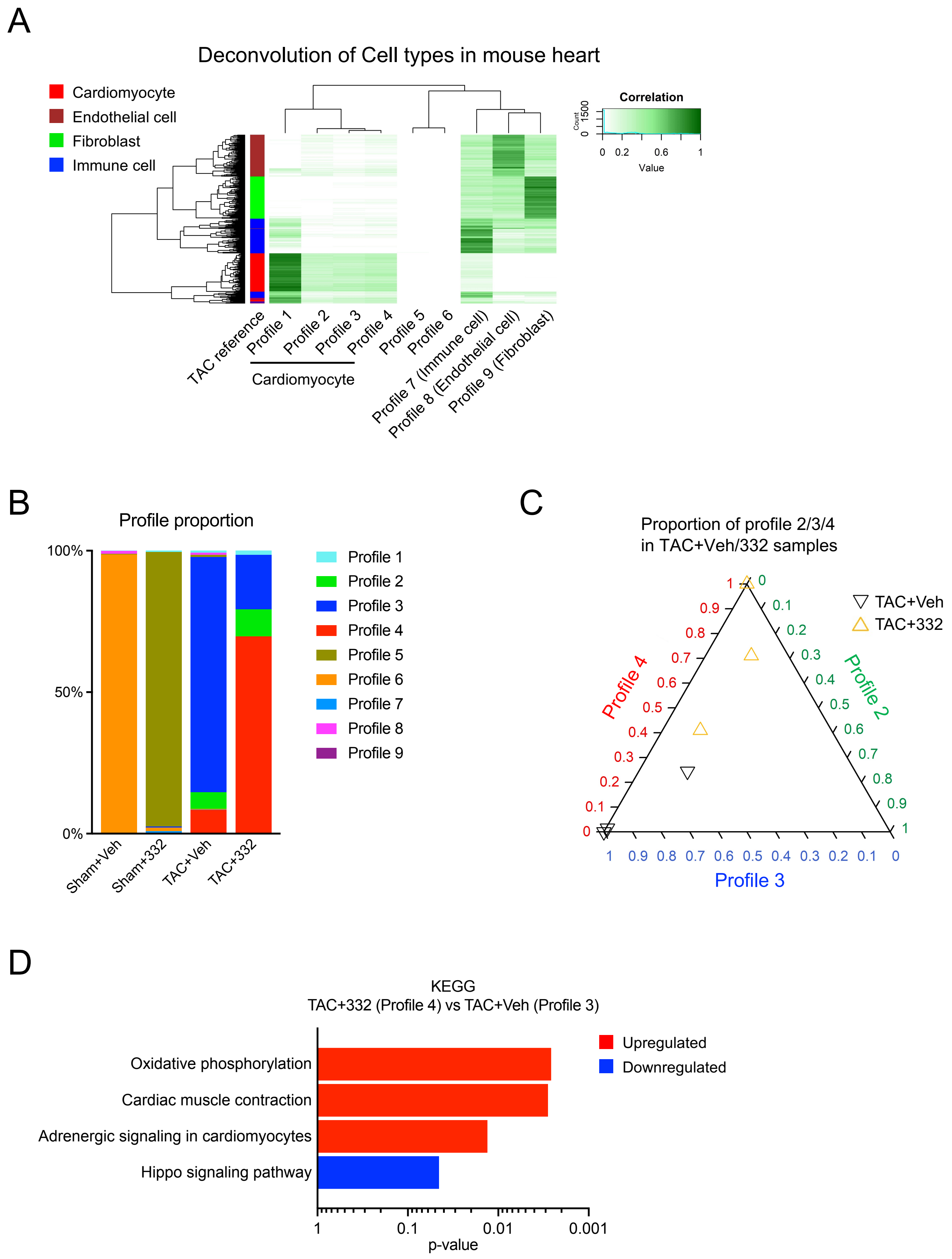

### Supplementary Figure 7

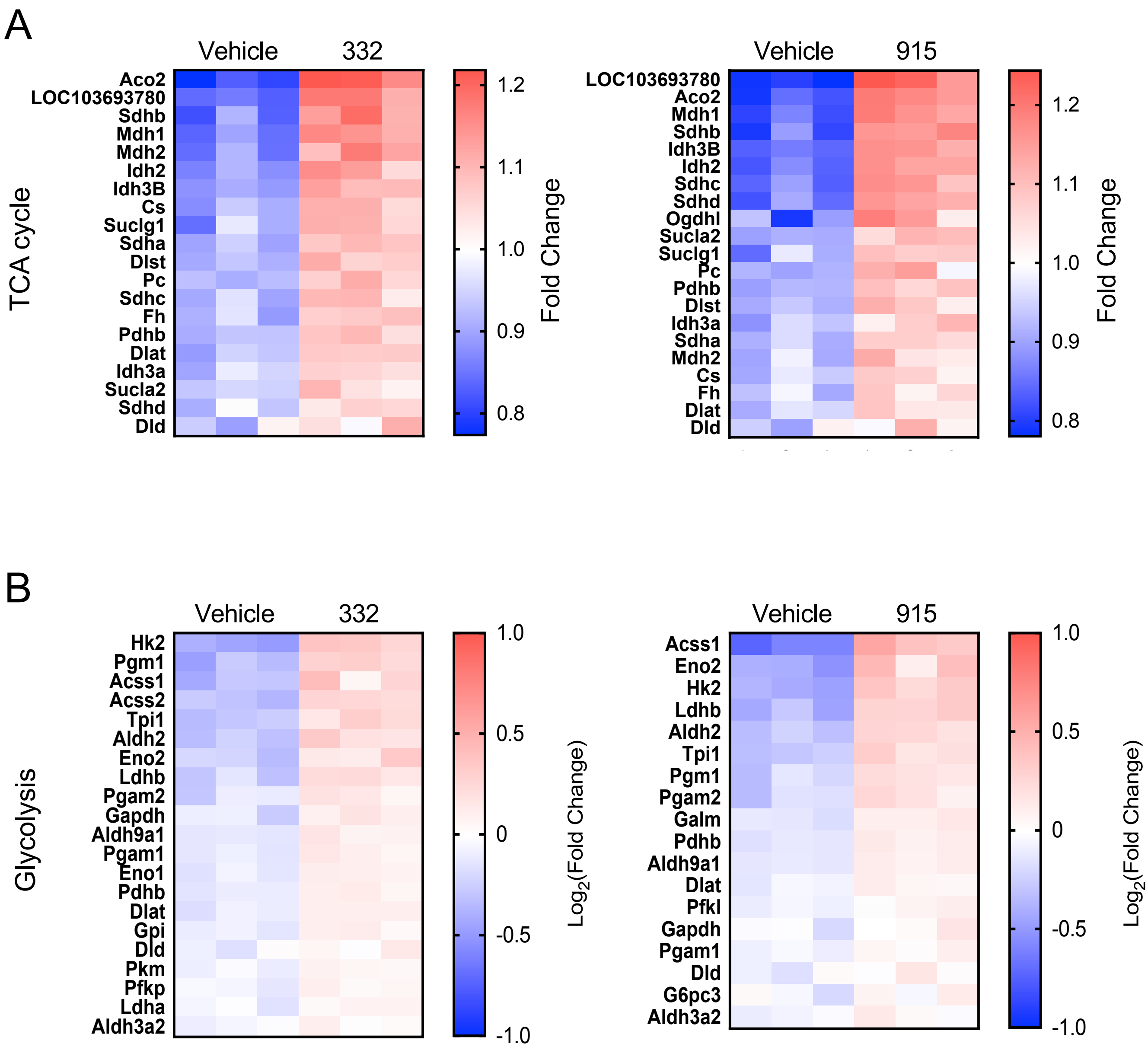

### Supplementary Figure 8

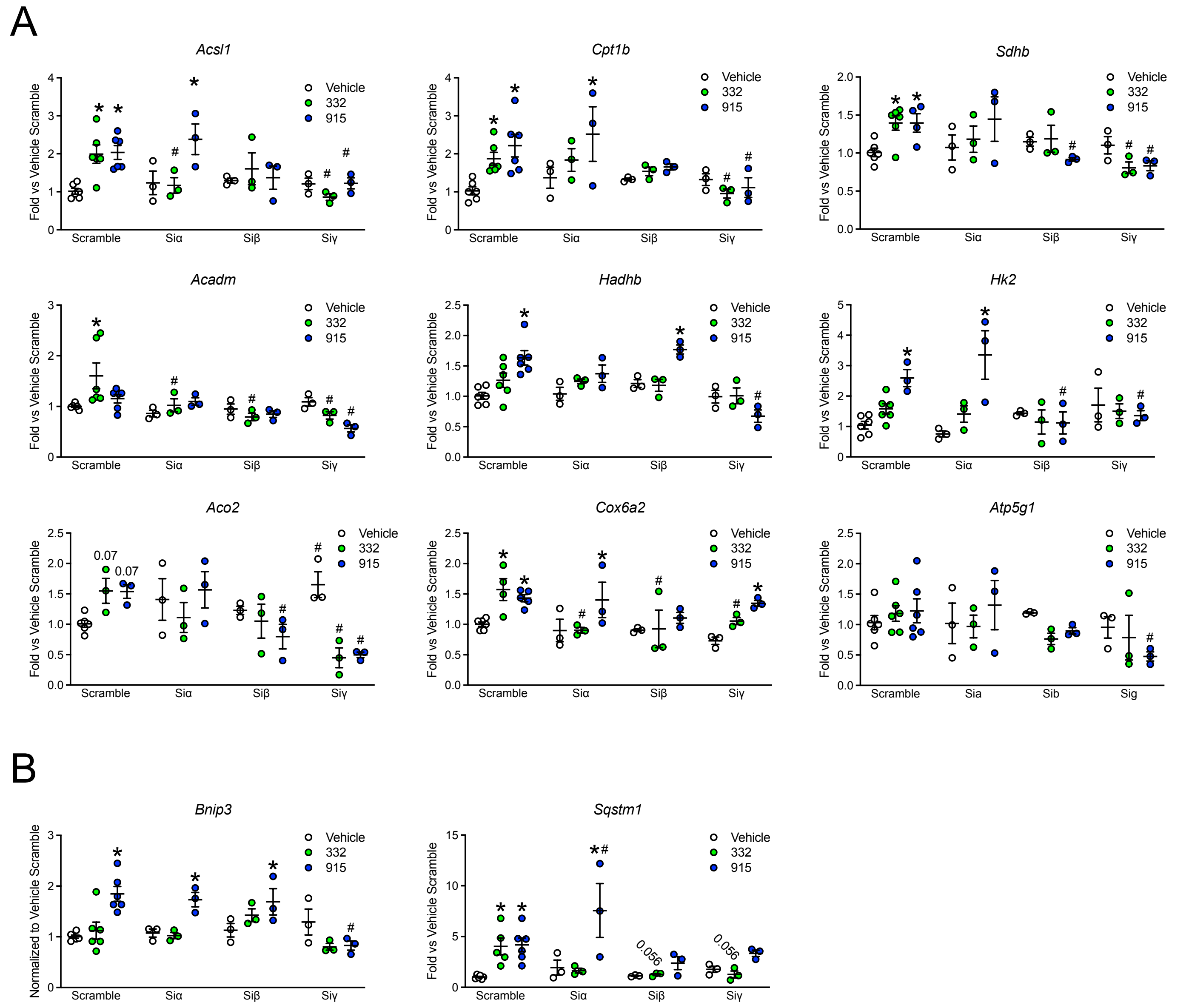

### Supplementary Figure 9

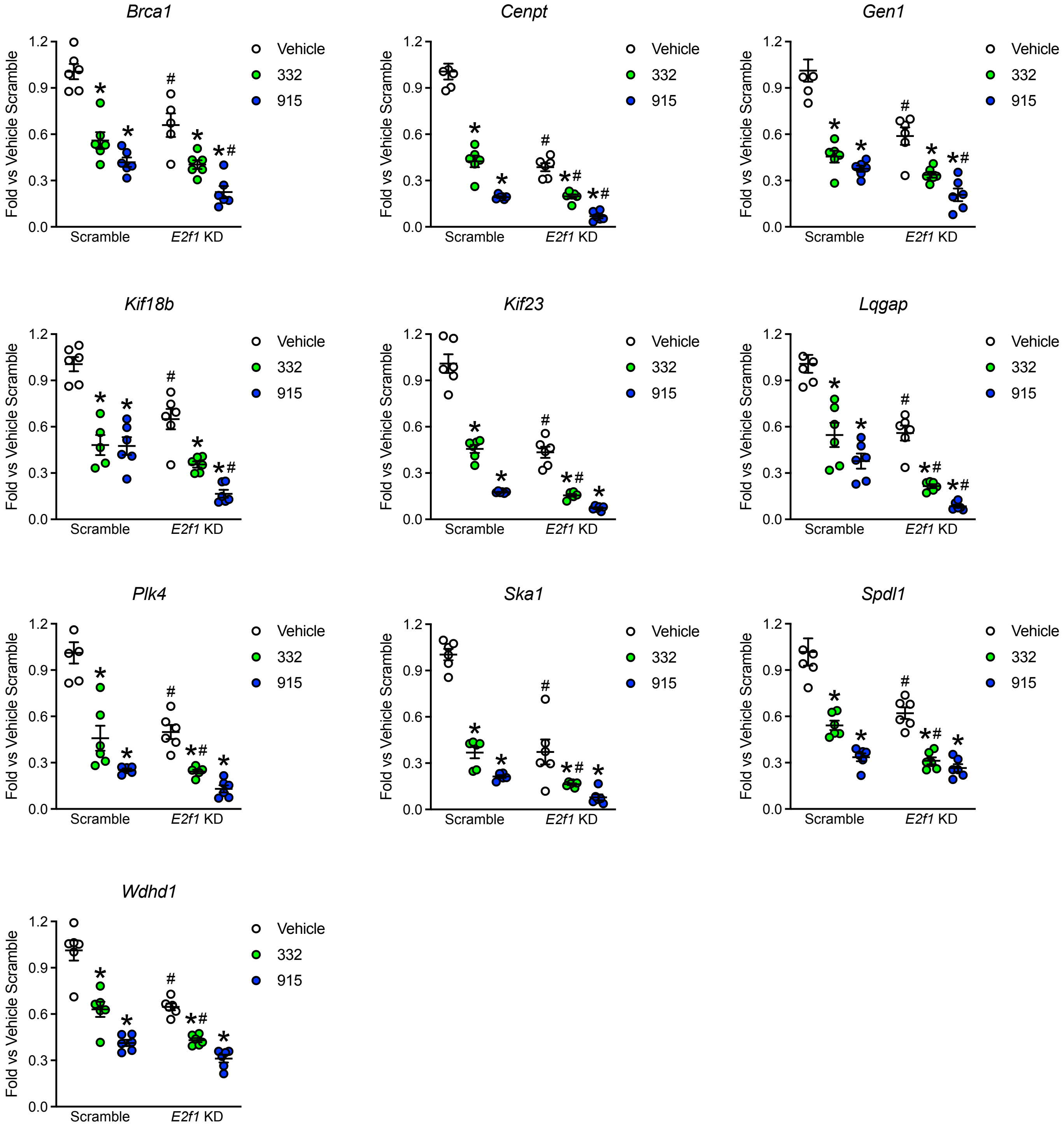

### Full unedited gel for Figure 3B

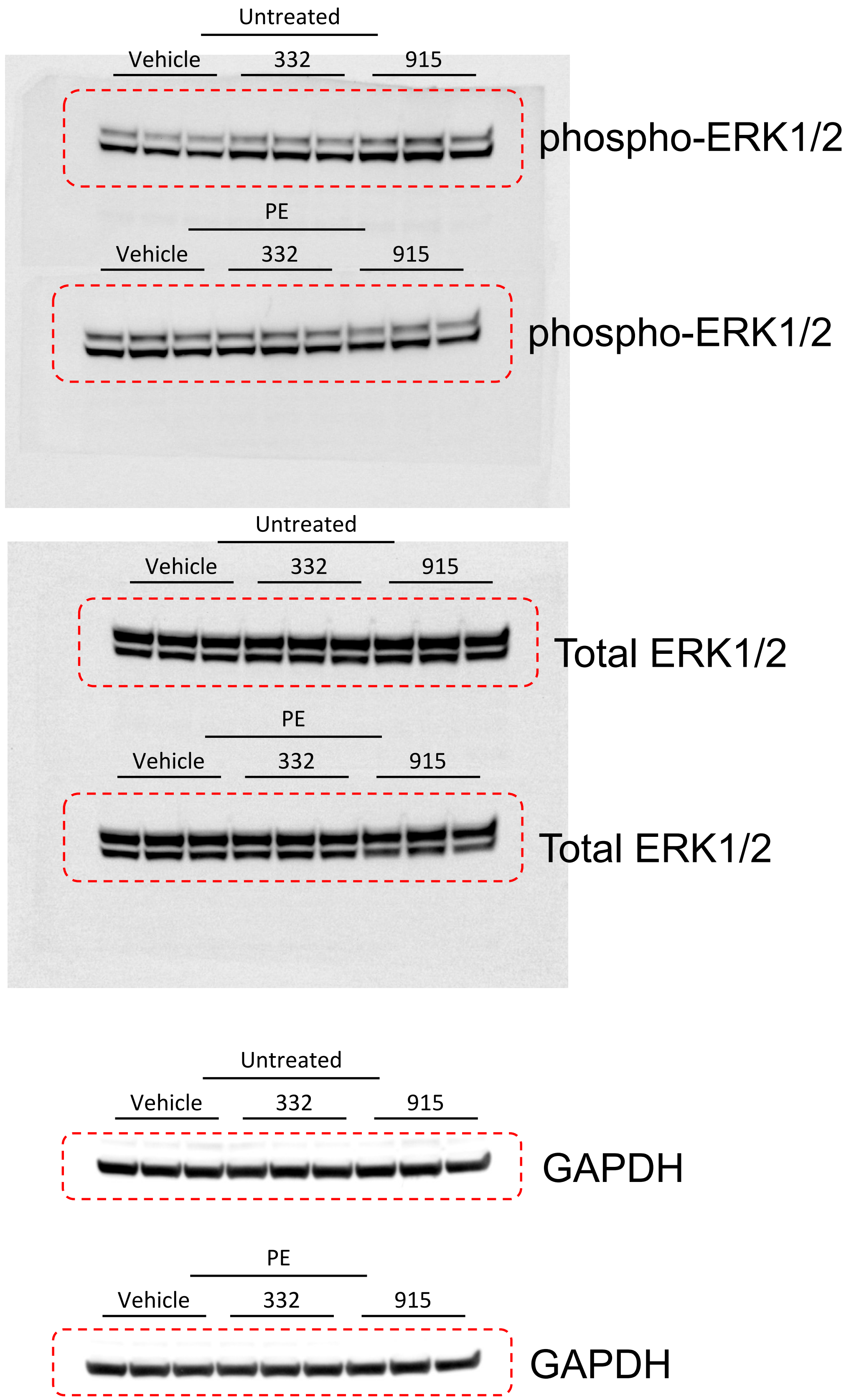

### Full unedited gel for Figure 6B

#### Experiment 1

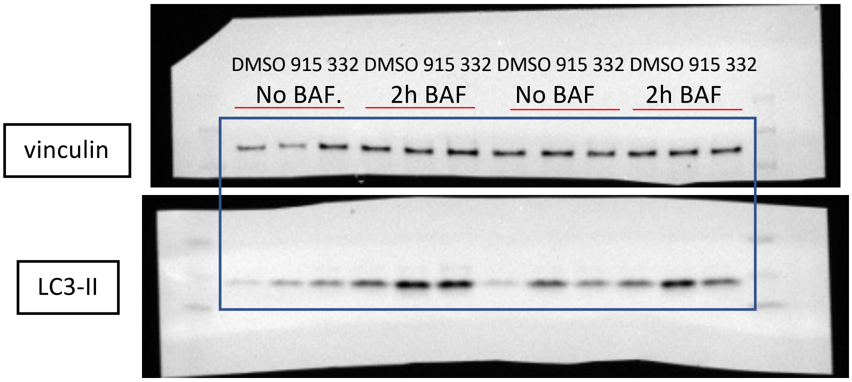


#### Experiment 2

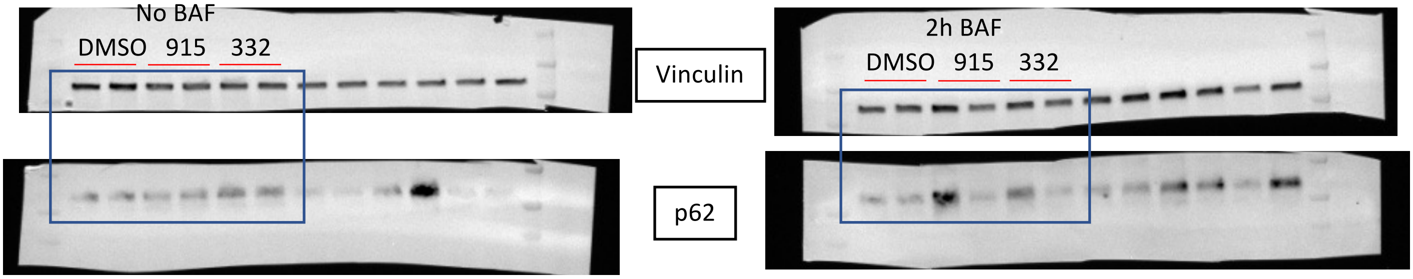


#### Experiment 3

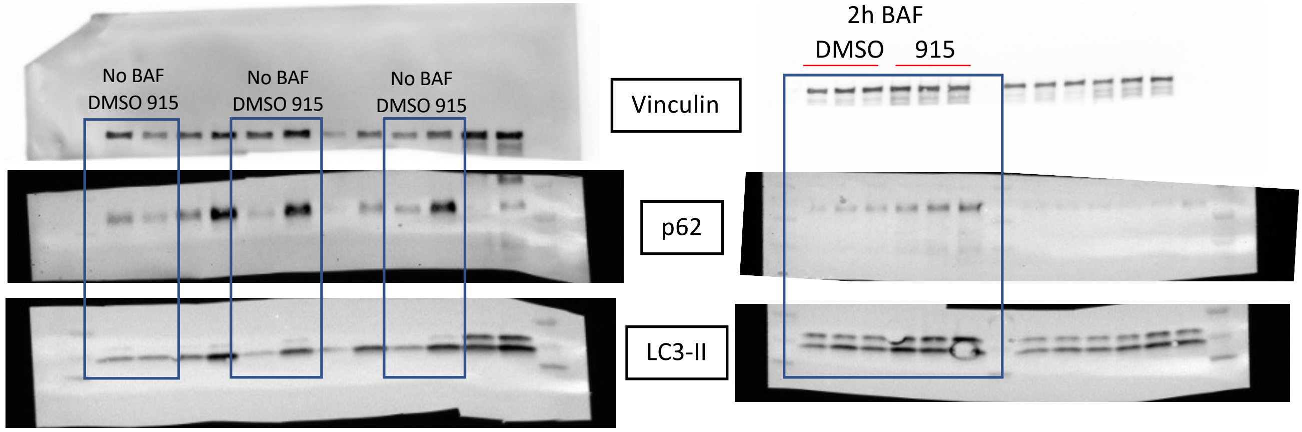


*bands not included in boxes are not used in analysis as they are part of a different experiment

Full unedited gel for Figure 7C

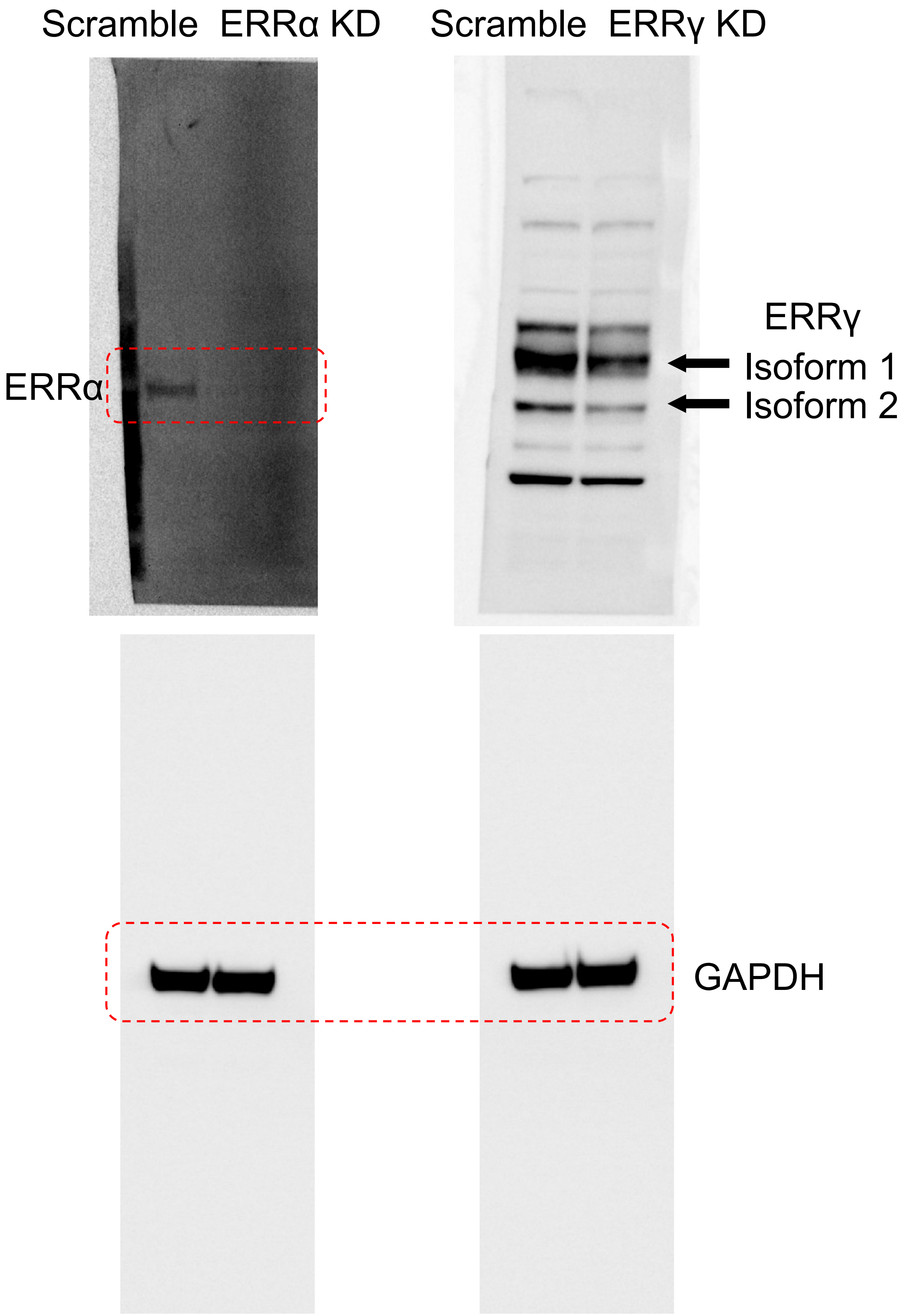
